# Predicting RNA-seq coverage from DNA sequence as a unifying model of gene regulation

**DOI:** 10.1101/2023.08.30.555582

**Authors:** Johannes Linder, Divyanshi Srivastava, Han Yuan, Vikram Agarwal, David R. Kelley

## Abstract

Sequence-based machine learning models trained on genome-scale biochemical assays improve our ability to interpret genetic variants by providing functional predictions describing their impact on the cis-regulatory code. Here, we introduce a new model, Borzoi, which learns to predict cell- and tissue-specific RNA-seq coverage from DNA sequence. Using statistics derived from Borzoi’s predicted coverage, we isolate and accurately score variant effects across multiple layers of regulation, including transcription, splicing, and polyadenylation. Evaluated on QTLs, Borzoi is competitive with, and often outperforms, state-of-the-art models trained on individual regulatory functions. By applying attribution methods to the derived statistics, we extract cis-regulatory patterns driving RNA expression and post-transcriptional regulation in normal tissues. The wide availability of RNA-seq data across species, conditions, and assays profiling specific aspects of regulation emphasizes the potential of this approach to decipher the mapping from DNA sequence to regulatory function.

## Introduction

A long-standing goal in genomics is to accurately assess the influence of each of the 3 billion nucleotides in the human genome with respect to gene-regulatory activity, ranging from chromatin accessibility and transcriptional activation to splicing and polyadenylation. A more illustrious goal is to predict how these regulatory functions change given genetic variation. Such predictions would dramatically improve researchers’ ability to interpret pathogenic mutations and prioritize functional variants at loci implicated in genome-wide association studies (GWAS), or even improve GWAS itself through functionally-informed discovery and fine-mapping [1, 2, 3].

Machine learning models trained to predict regulatory activity from DNA sequence have been quite successful at characterizing regulatory syntax and predicting genetic variant effects. Thus far, such models have focused on assays where measured activity is proportional to local sequencing read counts. For example, TF ChIP-seq or DNase/ATAC-seq aligned reads indicate a TF binding event or accessible DNA at the site where the reads align. This allows for accurate predictions using relatively short surrounding regions of sequence, typically 500-2, 000 bp [4, 5, 6, 7, 8, 9, 10].

In contrast, the most popular sequencing assay, RNA-seq, does not have this property; RNA-seq reads aligned across a transcript will depend on a much larger region of sequence containing the gene’s exons and relevant cis-regulatory elements. A read aligned to a gene’s 3^*′*^ end may be hundreds of thousands of nucleotides away from its promoter and enhancers that influence the magnitude of signal from the assay. Furthermore, RNA-seq coverage patterns integrate multiple layers of gene regulation – namely, transcription, splicing, termination/polyadenylation, and RNA stability. These properties make prediction of RNA-seq coverage from sequence challenging.

Previous models have only attempted to work with RNA-seq after its transformation to a gene expression matrix. By processing a large region centered on the transcription start site (TSS), several models can predict normalized gene counts [11, 12, 13]. This approach depends on accurate TSS annotation, suffers when multiple TSSs influence expression, and incompletely considers post-transcriptional regulation. Similarly, sequence-based models of splicing, polyadenylation, and RNA stability rely on transformed measurements extracted from RNA-seq data (such as percent spliced-in) that attempt to isolate these modes of regulation and make modeling tractable [14, 15, 16, 17, 18, 19, 20, 21, 22, 23]. However, such metrics inevitably struggle to describe complex splicing outcomes, de novo events, or the intricate and sometimes competitive relationship between transcription, splicing, and (intronic) polyadenylation [24, 25, 26].

Modeling RNA-seq coverage directly would have several benefits. First, RNA-seq is far richer than previously modeled assays. Although modeling multiple regulatory layers simultaneously is more challenging, it contains great promise; cross-talk between layers is common and their simultaneous consideration may improve models for each regulatory process. Second, current models for post-transcriptional regulation curate examples from genome annotations (for example, alternative spliced cassette exon junctions), which inevitably leads to loss of more complex examples. Training on genome-wide RNA-seq makes use of every relevant regulatory sequence instance, interpreting the coverage data in light of multiple processes. Third, there is a tremendous amount of RNA-seq data available, describing a wide variety of cell and tissue states across many species. Models trained on data from multiple species have been shown to outperform related models trained on single species [9], but chromatin profiling and the CAGE gene expression assays have been performed on far fewer species than RNA-seq.

Since mammalian genes often span hundreds of thousands of nucleotides, effective RNA-seq modeling requires working with very large sequences and algorithms that exchange information across large distances. Recent work on the Enformer model using self-attention demonstrates a path towards achieving this [13]. Thus, we set out to model RNA-seq and additional epigenetic assays’ coverage across diverse samples as a function of the underlying DNA sequence, without prior knowledge of gene annotation. We developed a data preprocessing pipeline, neural network architecture, and optimization strategy altogether named Borzoi that enabled effective learning of several layers of gene regulation in a single model. From a single RNA-seq experiment, Borzoi derives the primary cell type/state-specific TF motifs and a genome-wide map of nucleotide influence on gene structure and expression. Our model improved performance relative to Enformer on downstream tasks to identify distal enhancers and predict genetic variant effects on gene expression and introduced new capabilities to predict variant effects on splicing and polyadenylation that match or exceed state of the art. We anticipate this toolkit will accelerate progress to determine mechanisms by which the many unsolved human genetic associations affect traits.

## Results

### RNA-seq model design

RNA-seq is a nucleotide-resolution readout of transcribed and usually processed RNAs, precisely identifying, for example, splice junctions. Thus, modeling RNA-seq coverage at nucleotide-resolution would be ideal. However, the long span of mammalian genes means that we must also work with very long sequences to cover all exons and relevant regulatory elements. Computational limitations create a trade-off between these two considerations. We lean toward using longer sequences at the expense of some resolution, choosing 524 kb sequences for which we predict coverage in 32 bp bins.

Our neural network is illustrated in Figure 1A. We use the core Enformer architecture, which includes a tower of convolution- and subsampling blocks followed by a series of self-attention blocks operating at 128 bp resolution embedding vectors [27, 28]. Self-attention is a critical operation, allowing every pair of position vectors to exchange information. From this point, we make use of a U-net architecture to increase the resolution back to 32 bp [29, 30]. For each sequence length expansion (and resolution increase), we upsample the position vectors from the attention blocks and combine them with the corresponding feature map of equal size produced by the initial convolution tower (Methods). To go from embeddings representing 128 bp to those representing 32 bp, we perform this block twice, upsampling 2x each time. Continuing in our line of research naming biosequence ML models based on an analogy to talented scent-following hound dogs, we refer to models of this type for RNA-seq as Borzoi.

**Figure 1:**
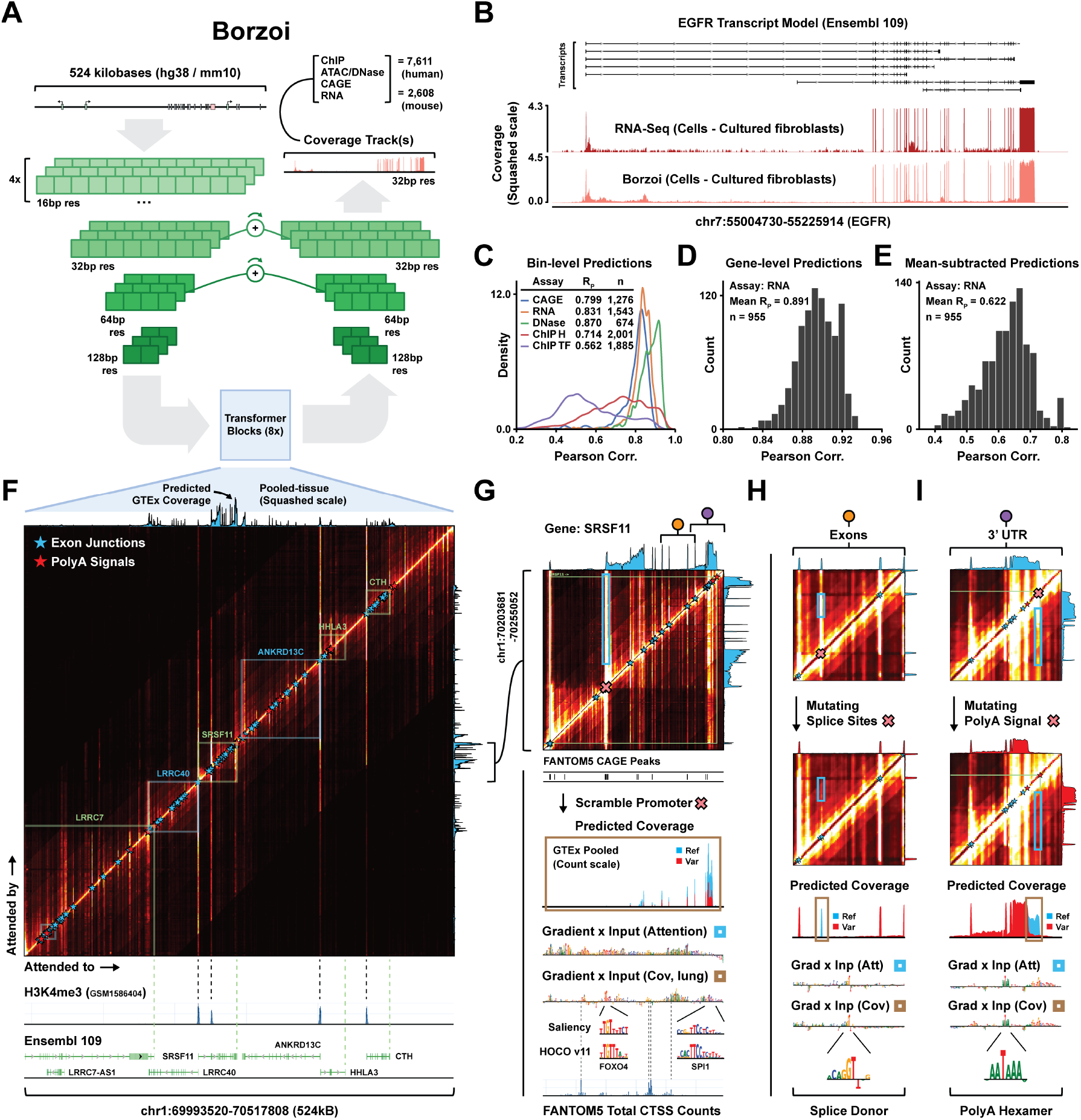
Borzoi - A neural network for predicting RNA-seq coverage from sequence. **(A)** The Borzoi neural network architecture consists of a number of convolution and downsampling layers, followed by a stack of self-attention layers with relative positional encodings operating at 128 bp resolution, similar to the Enformer architecture. The output is then upsampled through a number of deconvolutional layers with matched U-net connections to predict at 32 bp resolution. **(B)** RNA-seq coverage predictions for gene EGFR (GTEx tissue ‘Cells - Cultured Fibroblasts’). ‘Squashed scale’ refers to the transformed scale applied to the training data (Methods). **(C)** Pearson correlation on held-out test data (4-fold CV) across coverage tracks when predicting CAGE, RNA-seq, DNase or ChIP-seq (bin-level). **(D)** Gene-level Pearson correlation when comparing predicted to measured sum of RNA coverage across exons. **(E)** Gene-level Pearson correlation after quantile-normalizing the RNA coverage tracks and subtracting the average gene expression across tracks. **(F)** Attention weight matrix averaged across all 8 heads of the final transformer layers, shown for example region chr1:69993520-70517808. Average predicted RNA-seq coverage for 89 GTEx samples is shown above the attention heatmap. Ensembl transcript models and H3K4me3 tracks are shown below. **(G)**-**(I)** Enlarged view of the attention weight matrix for the SRSF11 gene, highlighting **(G)** a promoter region (and alternative TSS), **(H)** several introns and exons, and **(I)** the 3’ UTR. Gradient saliencies of either the output coverage tracks or the attention matrix (within the blue boxes) are displayed below each vignette. The regions highlighted in the saliency logos are either dinucleotide-shuffled (promoter) or mutated (exon and 3’ UTR) and the resulting coverage predictions are depicted above each logo (red). The altered attention matrices are also shown in **(H)** and **(I)**.

We chose to work with uniformly processed RNA-seq from the ENCODE project, providing 900 human and 600 mouse datasets [31, 32]. In addition, we included 2-3 replicates for each GTEx tissue processed by the recount3 project [33, 34, 35]. In order to help the model identify distal regulatory elements, we paired these data with the thousands of training datasets from the Enformer model, including CAGE, DNase, ATAC, and ChIP-seq tracks (Methods). To assess model performance variance and enable ensembling, we trained four replicate models with different held-out test sets.

### Borzoi accurately predicts RNA-seq and other sequencing assays

Despite the challenges of modeling RNA-seq coverage from only underlying DNA sequence, Borzoi predicts exon-intron coverage patterns with striking concordance for even long genes with many exons (Figure 1B). Test set predictions match RNA-seq coverage with a mean 0.83 Pearson R across human samples (Figure 1C). Performance is difficult to compare directly to Enformer due to differences in data processing (Methods). Nevertheless, test accuracies on the overlapping datasets are broadly similar, indicating competitive model training (Supp Figure S1A-D).

To study predictions at the gene-level, we sum coverage in bins overlapping exon annotations and compute log_2_. Transforming the experimental and predicted RNA-seq coverage similarly, we observe a mean 0.89 Pearson R across genes (Figure 1D). Finally, by quantile-normalizing the gene-level predictions and subtracting the average expression value taken over coverage tracks, we observe a mean 0.62 Pearson R (Figure 1E), indicating that the model explains a significant amount of the variation observed between tracks (such as tissue- and cell type-specific differences).

### Higher-level attention maps capture a comprehensive structure of genes

In order to impute RNA coverage, Borzoi must have learned a detailed representation of gene structure internally. We hypothesized that this learned gene structure is embedded in the higher-level attention layers. Figure 1F illustrates the average attention map of the two penultimate blocks for example locus chr1:69993520-70517808. The map highlights TSS regions, exon boundaries, and polyadenylation signals. Transcript elements tend to interact more within the same gene body, creating square-like structures that match annotated gene spans [36, 37]. The few positions attended to by many genes (seen as long vertical stripes) match peaks from H3K4me3 data [38]. In Supp Figure S1E, we extract the average attention magnitude of regions overlapping annotated H3K4me3 peaks, exons, and polyadenylation sites from held-out test loci and find a significant enrichment in intensity compared to matched background (*p <*1e-12).

We can visualize changes to global interactions in the attention map through in-silico sequence perturbation. Furthermore, while the attention map has 128 bp resolution, we can elucidate base-resolution determinants by computing the gradient saliency of attention in a region of the map with respect to the input sequence (‘Gradient x Input’; Methods). We can similarly compute the gradient of various statistics derived from the predicted RNA coverage pattern to focus on determinants of tissue-specific expression or post-transcriptional regulation. For example, Figure 1G shows the attention map of the SRSF11 gene. By dinucleotide-shuffling a 256 bp region overlapping the second TSS, the attention magnitude drops (Supp Figure S1F), as does predicted RNA coverage across the gene span. The gradients highlight TF binding motifs (e.g. putative FOXO- and SPI-like motifs) and promoter elements (e.g. SP motifs). Other attribution methods, e.g. In-silico Saturation Mutagenesis (‘ISM’), Window-shuffled ISM (‘ISM Shuffle’), and smoothed gradients integrated over categorical noise (‘Smoothgrad’) display roughly equivalent interpretations (Supp Figure S1G).

Attention gradients computed on smaller stripes within the gene body highlight sequences relating to splicing and polyadenylation. Intronic regions attend to the closest neighboring splice junctions (Figure 1H, Supp Figure S1H), suggesting that Borzoi is matching exon boundaries to infer low coverage across introns. When mutating the splice donor of one of the constitutive exons, attention drops along with RNA-seq coverage over the exon. Simultaneously, attention of the 3^*′*^ acceptor extends to the next viable 5^*′*^ donor site. Similarly, polyadenylation signals within SRSF11 attend to each other in order to adjust the coverage shape across the 3^*′*^ UTR. After mutating the distal site hexamer AATAAA, predicted RNA coverage drops upstream of the distal site and increases at proximal sites (Figure 1I, Supp Figure S1I). Proximal signals attend to the distal-most signal even for very long 3^*′*^ UTRs (20 kb) (Supp Figure S1J).

### Borzoi identifies regulatory motifs driving RNA expression in normal tissues

Borzoi was trained on a diverse set of RNA-seq samples and thus enables direct characterization of tissue-specific cis-regulatory drivers of expression by using attribution methods [39, 40, 41, 42, 43, 44]. We focused on a subset of 5 GTEx tissues to showcase this ability, namely whole blood, liver, brain, muscle and esophagus. We selected 1, 000 genes for each tissue with maximal TPM fold-change relative to other tissues and computed tissue-specific aggregated exon coverage gradients per gene. As an example, gradients at the position of maximal liver-specific saliency for gene CFHR2 highlight motif hits for CEBPA/B and HNF4A/G (Figure 2A).

**Figure 2:**
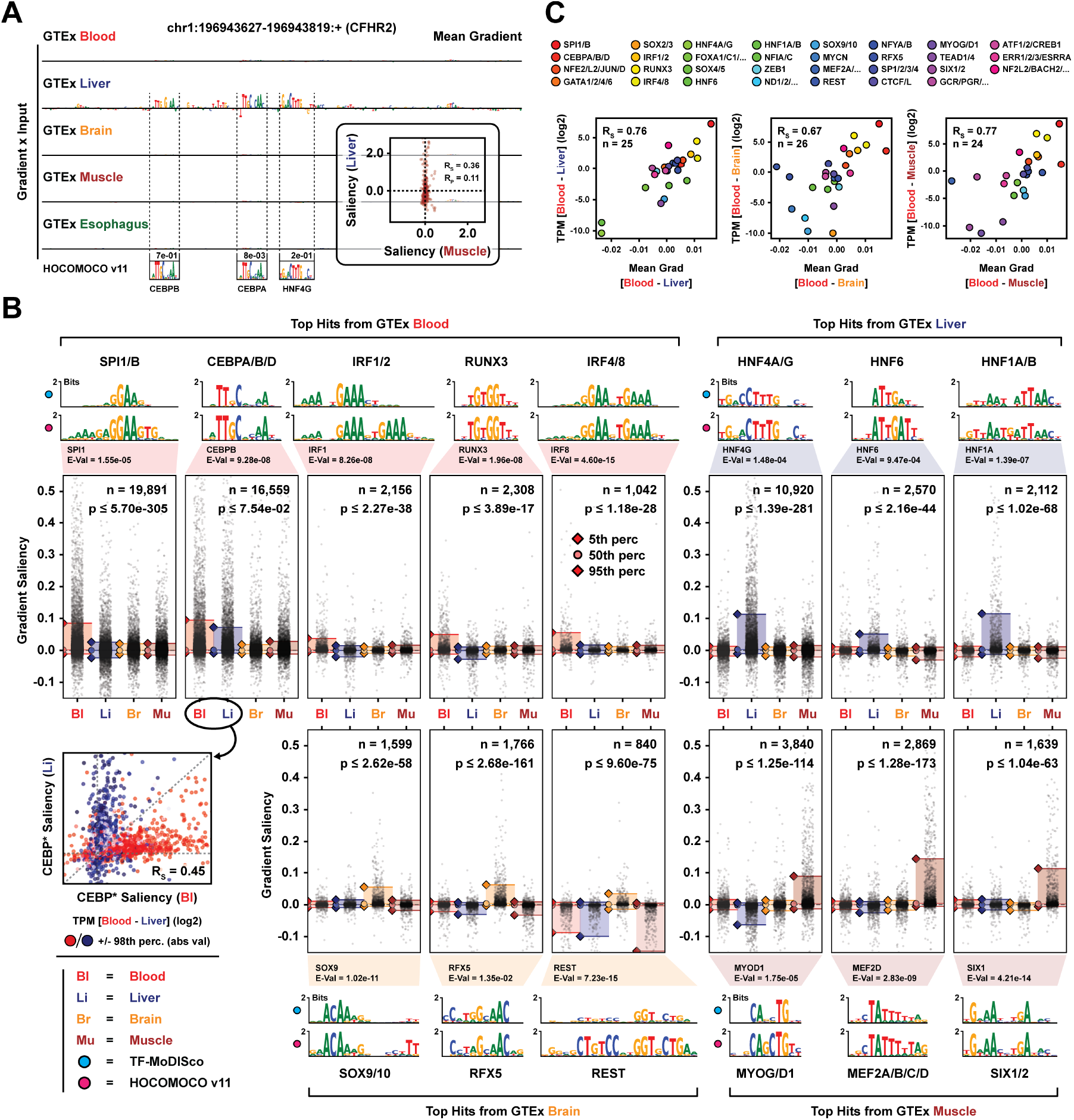
Identifying transcriptional cis-regulatory motifs through tissue-specific attribution. **(A)** Gradient attributions at the mode of maximum saliency for 5 GTEx tissues for liver-specific gene CFHR2. Likely motif hits and their PWMs from HOCOMOCO v11 are annotated. Inset: Comparison of nucleotide-level gradient saliencies for liver and muscle coverage tracks. **(B)** A selection of motif clusters identified by MoDISco from gradient saliencies corresponding to 4 GTEx tissues. Shown are the MoDISco PWMs, the best-matching PWMs from HOCOMOCO v11 and the distributions of tissue-specific gradient saliencies for seqlets belonging to a given cluster. P-values are computed using a two-sided Wilcoxon test between the tissue with largest absolute magnitude in gradient saliency (95th percentile) and the tissue with second largest saliency magnitude. Bottom left: Comparison of seqlet saliencies for putative CEBPA/B/D between the RNA coverage tracks of whole blood and liver. **(C)** Comparison between the average difference in gradient saliency of seqlets belonging to motif clusters for pairs of GTEx tissues and the difference in measured log-TPM for the corresponding TF genes. The median TPM of genes belonging to the same TF subfamily (HOCOMOCO v11) were averaged.

We first evaluated the quality of the gradient saliencies across the train / test set. Gradients computed from independent model replicates correlated reasonably well (Supp Figure S2A-B; median Spearman R = 0.53 between fold 0 and 1 for blood gradients in a 1, 024 bp window centered at the mode of importance), as did gradients and ISM maps (Supp Figure S2C; Spearman R = 0.72 within a 192 bp window around the mode of importance). When evaluating model fold 0 on a subset of held-out test genes, its tissue-specific differential coverage predictions correlated with measured TPM fold changes as well as for trained-on genes (Supp Figure S2D; median Spearman R = 0.82 vs 0.87 for train / test). The same trend was observed when comparing the average difference in gradient saliency to TPM fold changes (R= 0.64 for both train and test). All downstream analyses rely on the average gradient across the 4 model replicates.

Next, for each set of 1, 000 genes chosen for a specific tissue, we selected the corresponding gradients for that tissue and subtracted the average gradient of all other tissues, obtaining residual scores focused on tissue-specific regulation.

We ran TF-MoDISco, a de novo motif clustering tool [45], on each set of 1, 000 genes for all 5 tissues. The MoDISco clusters were matched to their most likely motif using the Tomtom MEME suite and HOCOMOCO v11 [46, 47] and the identified seqlets were pooled across MoDISco reports by motif match. A selection of top-scoring motifs are shown for whole blood, brain, liver and muscle alongside their gradient saliency distributions in Figure 2B (see also Supp Figure S2E-F). We detect many well-known regulatory motifs for each tissue, such as SPI1/B and IRF4/8 (for blood), HNF4A/G and HNF1A (for liver), SOX9 and REST (for brain), and MYOD1 and MEF2D (for muscle). For motifs with significant gradient saliency in multiple tissues, we generally find that these saliencies originate from distinct loci (Figure 2B, inset). We only identify a few weakly tissue-specific motifs for esophagus (Supp Figure S2G), such as TP53, COT2 and SRF. While we do find GATA motif hits for the GTEx blood gradients, we were curious as to why their saliency scores were relatively small in comparison to other drivers of blood-specificity. Re-running the analysis using gradients derived from K562 RNA-seq tracks, we find GATA to be one of the highest-scoring clusters with significantly larger saliencies in K562 compared to whole blood (Supp Figure S2H), suggesting that GATA is a significant driver of expression in the erythroleukemia cell line K562 but less so in a diverse mixture of blood. Directly comparing the measured expression quantiles of TF genes in GTEx whole blood and K562 experiments supported our inferences; we observed a considerable upregulation of SPI1, CEBPB and IRF motifs in whole blood alongside a decrease in GATA1/2 TPM quantiles relative to K562 (Supp Figure S2I).

Finally, we aggregated the difference in gradient saliency for each pair of tissues among seqlets matching each TF, obtaining a scalar score that describes the importance of a particular TF in one tissue relative to another. When compared to observed TPM fold changes for the TFs in the corresponding tissues (averaged across TFs of the same HOCOMOCO subfamily), we observed a strong concordance across all tissue pairs (Figure 2C and Supp Figure S2J-K). For example, Spearman R reached 0.77 when comparing TF saliency in blood and muscle. Note that a repressor element such as REST should be off-diagonal in comparisons to brain, so we do not expect a perfect correlation. Altogether, Borzoi is capable of producing detailed gene regulatory sequence maps that highlight specific TF regulators.

### Improved utilization of distal regulatory sequence features for gene expression prediction

Although promoter regions explain a substantial proportion of the variance in expression across genes [11], distal regulatory elements like enhancers are critical to cell and tissue-specific regulation [49, 50, 51, 52]. Though imperfect, Enformer competitively ranks distal regulatory elements for their gene-specific enhancer activity, as validated by CRISPR screens [13, 53, 54, 55, 56, 57, 58]. We applied a similar procedure to score putative enhancers with Borzoi. For each target gene, we computed input gradients of the aggregated exon coverage prediction in RNA-seq samples with matched cell type (K562) and smoothed the scores in a local window using a Gaussian filter. Compared to Enformer, we can score sites that are up to twice as far away from the gene, 262 kb, and we make use of exon-, rather than TSS-annotations, which are generally more robust to alternative isoforms. Figure 3A displays the gradient attributions for example gene HBE1, where Borzoi correctly identifies a group of proximal enhancers (distance to TSS *<* 20, 000 bp). Figure 3B shows attributions for another example (MYC), where Borzoi identifies a distant enhancer located more than 200, 000 bp away, though false positives are also present.

**Figure 3:**
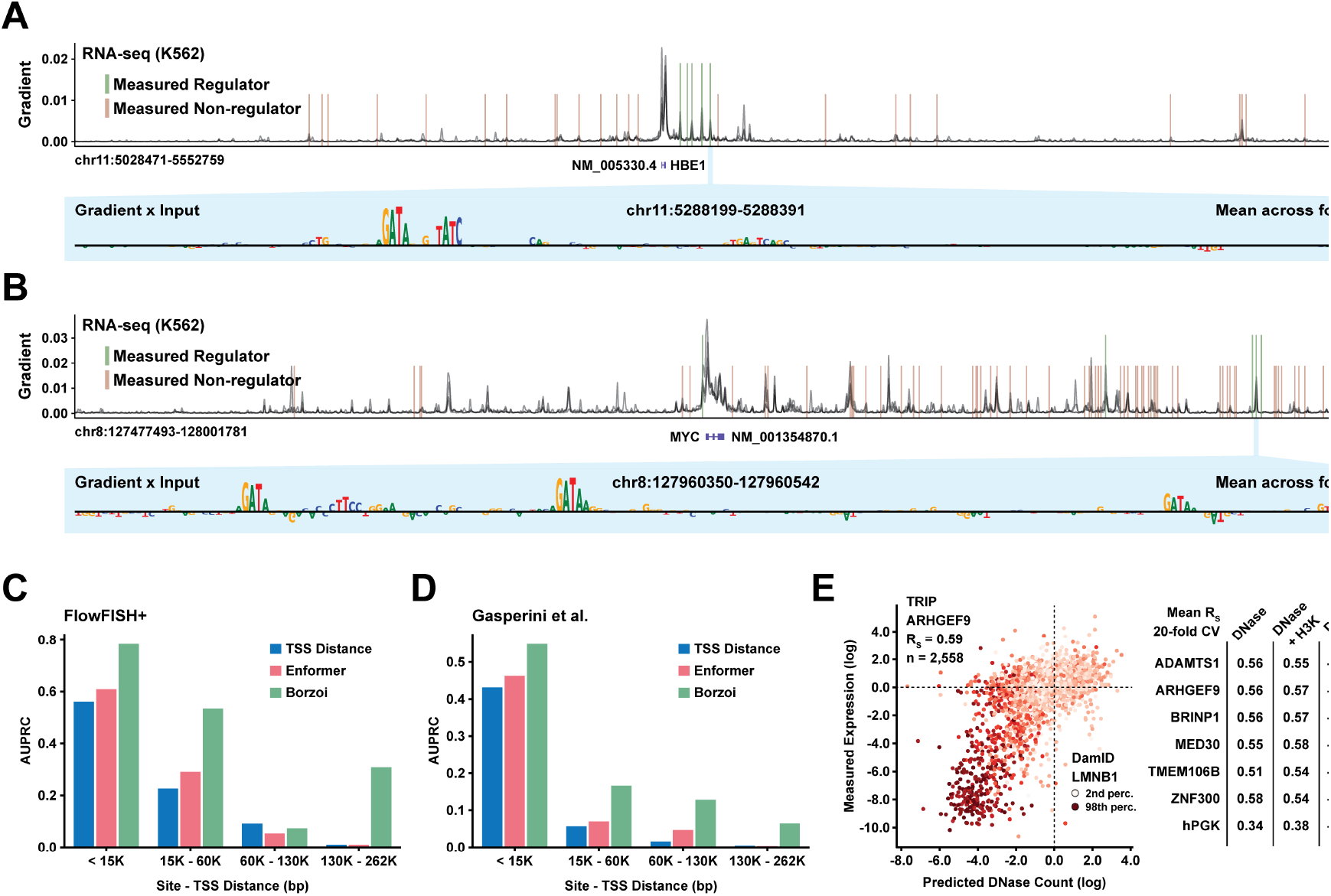
Predicting the impact of context and distal regulatory elements on gene expression. **(A)** Aggregated exon coverage gradient saliency for HBE1 across the 524 kb input (curves for 4 model replicates are shown). CRE regions that have been measured to regulate (green) or not regulate (red) HBE1 are annotated [48]. Input-gated gradients for a 192 bp window centered on the most distal enhancer is displayed at the bottom. **(B)** Exon coverage gradients for MYC. **(C)** Average precision (AUPRC) when using a statistic computed from the Borzoi or Enformer gradient saliencies within a local window around each CRE locus to classify whether it regulates the target gene (measurements from Nasser et al., 2021). **(D)** AUPRCs when using the Borzoi or Enformer gradient scores to classify regulating / non-regulating CRE loci in data from Gasperini et al. (2019). **(E)** Left: Predicted vs measured expression levels of TRIP reporter constructs based on Borzoi DNase coverage in K562 (Promoter: ARHGEF9). Color = DamID LMNB1 measurements. Right: Average Spearman R (20-fold CV) when predicting TRIP expression based on different scores.

When comparing Borzoi, Enformer and a distance-to-TSS baseline on their ability to classify measured positive from negative enhancer-gene interactions in data from Fulco et al. (2016, 2019), Klann et al. (2017) and others [55, 57, 58, 59, 60, 48], we find that Borzoi has superior AUPRC in all distance categories but one and highest AUROC in all categories (Figure 3C; Supp Figure S3A). For the larger dataset from Gasperini et al. (2019) [53], Borzoi has the highest AUPRC / AUROC at all distances (Figure 3D; Supp Figure S3B). At a distance of 130 - 262 kb from the target gene, Borzoi has more than 15x higher AUPRC than the TSS baseline.

To further demonstrate the model’s reliance on broader genomic context for its predictions, we analyzed expression data of 7 distinct promoters that had been integrated into thousands of genomic positions by the TRIP assay [61, 62]. We predicted activity scores from multiple classes of coverage tracks, including DNase, histone modifications, CAGE and RNA-seq. In general, the scores derived from DNase tracks were most concordant with the measured expression levels (Figure 3E, Supp Figure S3C; 20-fold CV Spearman R = 0.56 for promoter ARHGEF9) and these predictions were better correlated with measured expression than LMNB1 DamID-seq, which measures nuclear lamina interactions where expression tends to be lower (e.g. Spearman R = −0.39 for ARHGEF9). We speculate that the lower performance of the RNA-seq tracks compared to DNase is a result of the reporter construct being dissimilar from training examples of genes in the genome (i.e. no introns, PiggyBac transposons, etc.).

### Borzoi predicts the impact of genetic variation on gene expression

Accurately predicting the influence of genetic variant alleles on gene expression is a major need of the genome research community in order to determine the regulatory mechanisms of genetic associations in human populations. Here we evaluated Borzoi’s ability to distinguish fine-mapped GTEx eQTLs from a set of matched negatives, controlling for TSS distance [1]. As an example, Figure 4A (and Supp Figure S4A) shows the predicted RNA coverage track and sequence attributions for SHTN1 in GTEx ‘Whole Blood’ when inducing SNP rs1905542, alongside measured whole blood coverage in tissues from GTEx individuals harboring each allele. Borzoi correctly predicts upregulation of SHTN1 expression due to creation of a CEBP binding motif with predicted epistatic interactions to nearby blood-specific motifs (IRF1/2, SPI1/B) [63, 64, 65, 66]. Additional eQTL examples are visualized in Supp Figure S4B-C.

**Figure 4:**
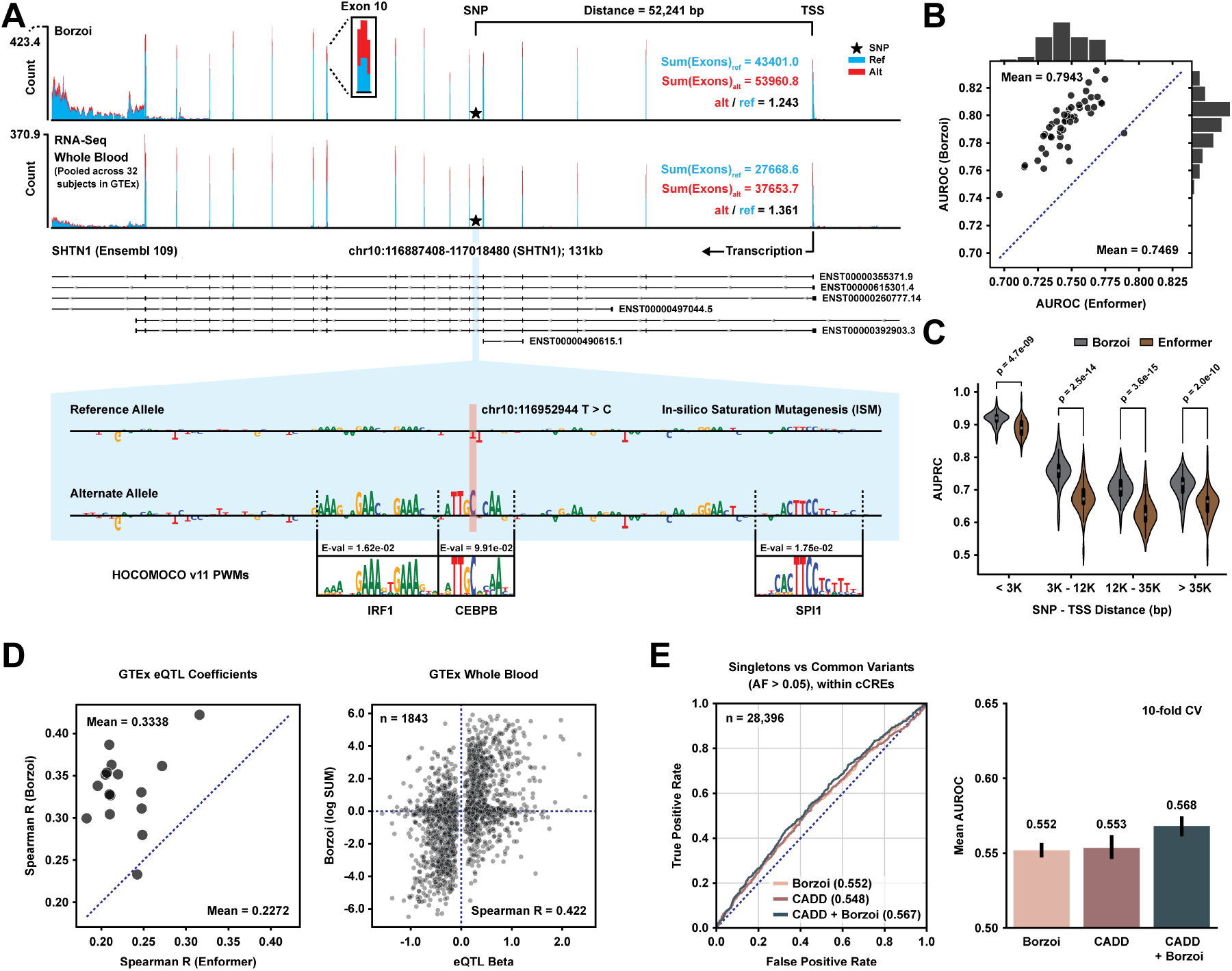
Borzoi predictions of variant effects align with eQTL results and negative selection. **(A)** Example eQTL rs1905542. Shown are the predicted RNA-seq ‘Whole Blood’ coverage tracks from GTEx for the reference (blue) and alternate (red) alleles, as well as the measured, aggregated RNA-seq coverage in ‘Whole Blood’ for 32 homozygous carriers of the reference allele and 32 hetero- or homozygous carriers of the alternate allele. ISM maps are shown (with equally scaled y-axes) at the bottom along with likely motif hits. **(B)** AUROC per GTEx tissue when classifying fine-mapped causal eQTLs from distance-matched negatives. Results are compared between Borzoi and Enformer. **(C)** Comparison of tissue-specific GTEx eQTL classification performance as a function of distance to the TSS. P-values are computed using a two-sided Wilcoxon test. **(D)** Left: Comparison of Spearman correlation coefficients between predicted and observed eQTL effect sizes from GTEx, using either Borzoi or Enformer with the differential log-sum coverage statistic (‘SUM’; Methods). Right: Predicted vs observed eQTL effect sizes in GTEx Whole Blood based on SUM scores derived from Borzoi’s RNA-seq tracks. **(E)** Precision vs recall when classifying singleton variants from common variation (AF *>* 0.05) from gnomAD. Average AUROCs are shown to the right (10-fold CV). All variants were sampled from ENCODE candidate cis-regulatory elements. AUROC scores displayed in legend.

Similar to Enformer, Borzoi predicts coverage across a large sequence region for many experimental tracks from which a variant effect score must be distilled. Briefly, for RNA-seq tracks we compute either the log-sum fold change or L2 norm of differential coverage across exons (Methods). Using Borzoi with an L2 score was superior to Enformer and its original score aggregation (sum) at discriminating eQTLs (Figure 4B-C), with a mean AUROC of 0.794 per tissue compared to Enformer’s AUROC of 0.747. Borzoi still outperformed Enformer when using a single model instead of the ensemble of four (AUROC = 0.788) or when switching to the original sum statistic (AUROC = 0.772). Enformer with an L2 score partially closes the gap (AUROC = 0.770). Limited to the same tracks as Enformer (ChIP, DNase, ATAC, and CAGE), Borzoi already achieves greater accuracy (AUROC = 0.784). Limiting to only the RNA-seq tracks achieves a similar 0.785 AUROC. Beyond inferring whether a SNP will have a significant effect on expression, there has been shifted focus recently towards imputing personalized gene expression values [67, 68]. This problem demands sign concordance and well-calibrated predictions. Borzoi exhibits greater Spearman correlation than Enformer when comparing effect size predictions to fine-mapped eQTL coefficients (Figure 4D). For example, for blood, Borzoi has a Spearman R of 0.422. Model ensembling accounts for a part of the performance increase (mean *R* = 0.334 across tissues), but Borzoi outperforms Enformer with even a single model (mean *R* = 0.292 compared to *R* = 0.227).

We hypothesized that Borzoi could be used to prioritize the affected eGene from the set of genes surrounding an eQTL association. As a baseline, we predicted the nearest gene to be the eGene, where distance between an eQTL and a gene is defined as the average inverse distance to an annotated TSS across all variants within the fine-mapped credible set, weighted by each variant’s posterior causal probability (PP). Using Borzoi, we aggregated L2 scores for all variants within the eQTL’s credible set, weighted by their fine-mapping PP and ranked the genes by their maximum aggregated score. Borzoi prioritized the true eGene at a similar rate as TSS distance across GTEx tissues (mean accuracy 63.4% vs 62.7%, Supp Figure S4D), suggesting that the model has not learned complex determinants of enhancer specificity.

To further test the utility of Borzoi-derived variant scores, we investigated the degree to which the model can distinguish common variation, which is nearly always benign, from singletons, which are relatively enriched for pathogenicity, in the GnomAD database [69, 70]. For each singleton sampled, we sampled a common variant, controlling for trinucleotide context and alternative allele. For comparison, we considered CADD (v1.6) scores [71, 72], which are derived from multiple sources including sequence conservation and functional annotations. While both methods have limited discriminative power on these data (which is expected due to the clear but subdued relationship between function and allele frequency), Borzoi outperformed CADD (mean AUROC 0.54 vs 0.53, Supp Figure S4E). Restricted to ENCODE candidate cis-regulatory elements, Borzoi and CADD performed equally well (mean AUROC 0.55). Combining their scores resulted in the highest accuracy (mean AUROC 0.57, Figure 4E). Borzoi’s ability to learn the sequence drivers of multiple modes of regulation thus make it a useful candidate for prioritizing variants impacting diverse regulatory processes and augmenting existing CADD annotations.

### Inference of 3^*′*^ UTR APA isoforms from RNA coverage

The 3^*′*^ untranslated region (3^*′*^ UTR) harbors a multi-layered cis-regulatory code that jointly controls 3^*′*^-end formation and transcript stability. When the mRNA is transcribed, short regulatory regions called polyadenylation signals (PASs) recruit the 3^*′*^-end processing machinery which terminates transcription and polyadenylates the transcript [73]. In the case of multiple PASs within the same gene, competition for cleavage and polyadenylation creates isoforms with distinct 3^*′*^ ends (Alternative Polyadenylation, or APA). The strength of a PAS is regulated by binding motifs for core polyadenylation regulators such as CFIm, CstF and CPSF as well as auxiliary proteins such as SRSF- and HNRNP proteins [74, 75]. While this code is local to the 3^*′*^ cleavage site, the competition created between polyadenylation signals and splice sites creates a non-linear and long-ranging cis-regulatory code.

We reasoned that Borzoi has learned these layers of post-transcriptional regulation in order to predict RNA coverage at transcript ends. Figure 5A shows predicted and measured tissue-pooled coverage for an example locus in the 3^*′*^ UTR of SRSF11. Motifs for well-known polyadenylation regulators (CFIm, CPSF, CstF, etc.) emerge from computing a ratiometric input attribution score to delineate the sequence elements driving coverage at the distal-most cleavage site relative to coverage elsewhere. When comparing the predicted distal-to-proximal polyadenylation coverage ratios of held-out genes to measurements in GTEx samples, Borzoi had an average Spearman R of 0.88 across folds (Figure 5B). These predictions also correlated with measured isoform abundances in PolyADB v3 [76, 77] (Figure 5C; average Spearman R = 0.64). Finally, by computing gradient saliencies of polyadenylation coverage ratios for genes from the Gasperini set [53] and clustering the gradients using TF-MoDISco [45], we recapitulated the core 3^*′*^-end processing motifs as top hits (CFIm, CstF, Elavl) (Figure 5D).

**Figure 5:**
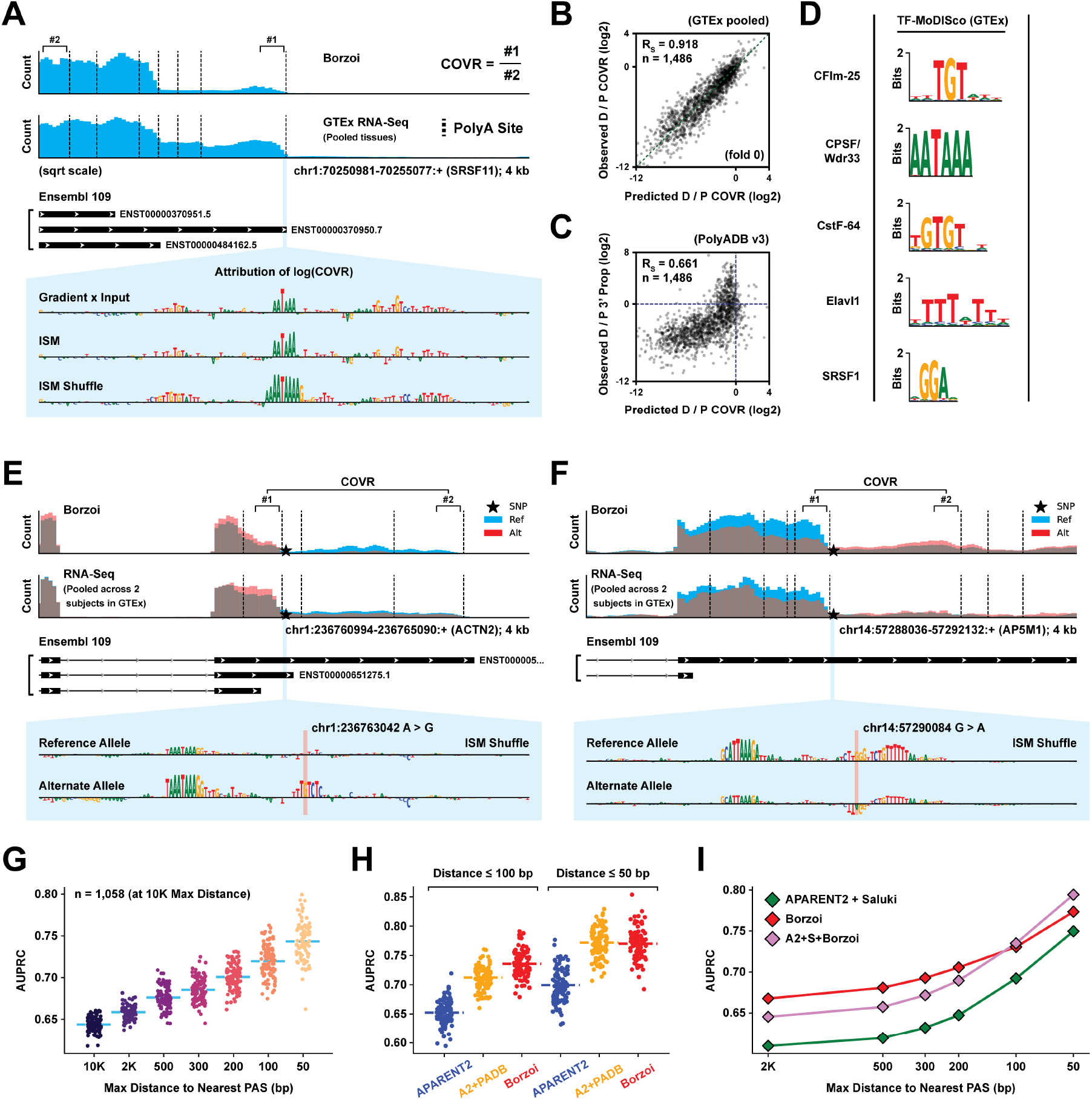
Predicting Alternative Polyadenylation and 3’ Polyadenylation QTLs. **(A)** Example locus (distal PAS of SRSF11 gene). Shown are the predicted and measured pooled coverage from GTEx samples (plotted in square-root scale due to large expression values). Calculation of polyadenylation-centric coverage ratios (COVR) is illustrated in the figure. Attribution scores (gradient saliency, ISM and ISM Shuffle) at the bottom. **(B)** Predicted vs measured coverage ratio between the distal-most and proximal-most PAS of each gene in held-out test data (tissue-pooled GTEx). **(C)** Predicted RNA-seq coverage ratio vs measured isoform proportions (PolyADB). **(D)** MoDISco PWMs of well-known APA regulators, obtained from pooled GTEx coverage ratio gradients. **(E)** Predicted RNA-seq coverage (GTEx pooled) for variant rs114880747, along with measured coverage in 2 individuals with the reference allele and 2 heterozygous individuals (3 tissues). Attribution scores (bottom) suggest gain of a CstF motif. **(F)** Predicted and measured coverage in individuals without and with variant rs80168986 (2 individuals, 3 tissues each). Attribution scores (bottom) suggest gain of a HNRNPA1 motif. **(G)** AUPRC when classifying fine-mapped paQTLs from GTEx based on predicted RNA-seq coverage ratio statistics (tissue-pooled GTEx), plotted as a function of decreasing distance threshold to the nearest 3’ UTR PAS. Each dot represents a permutation test (Methods). **(H)** paQTL classification AUPRC comparing variant effect predictions of Borzoi, APARENT2 and APARENT2+PolyADB. **(I)** paQTL classification AUPRC as a function of decreasing distance threshold to the nearest PAS. ‘A2+S+Borzoi’ represents an ensemble of all models.

We generally do not find concordance between 3^*′*^ UTR attributions from Borzoi and Saluki, which is a sequence-based model of mRNA half-life [23], nor to mutagenesis experiments [78]. It is however hard to conclude that Borzoi has not learned determinants of transcript stability, since those motifs overlap with polyadenylation elements (e.g. ARE-like motifs which are salient downstream motifs, and CFIm which binds to TGTA[A/T]). In Supp Figure S5A, we do observe correlation between the codon-aggregated gradient saliencies of gene exon coverage and MPRA measurements from Forrest et al (2020) [79] (Pearson R = 0.59), which suggests the model has learned codon contribution to stability.

### Functional polyadenylation variant effect prediction

Another important class of disease variants modulate expression by altering 3^*′*^ mRNA processing [80]. We next investigated Borzoi’s ability to score 3^*′*^ UTR variants that impact mRNA isoform abundance. We curated fine-mapped 3^*′*^ QTLs from the eQTL Catalog [81, 82] (paQTLs; n = 1, 058) and constructed a set of expression-matched negatives, controlling for distance to the nearest 3^*′*^ cleavage site. We calculated variant effect scores as the maximal absolute change in predicted coverage ratio between any 3^*′*^ cleavage junction from tissue-pooled GTEx tracks. An example paQTL is shown in Figure 5E (and Supp Figure S5B), where an A*>*G change results in gain of a putative CstF binding motif (rs114880747). Compared to RNA-seq tracks of GTEx individuals harboring the alternative allele, Borzoi correctly predicts upregulated site usage and the input attributions highlight the motif gain. Conversely, Figure 5F (and Supp Figure S5C) highlights a G*>*A change (rs80168986) that results in decreased polyadenylation due to the creation of an HNRNPA1 site. More examples are highlighted in Supp Figure S5D.

The variant effect scores derived from the predicted RNA-seq tracks discriminated paQTLs from the matched negatives with a monotonic increase in accuracy at closer distances to the nearest PAS (Figure 5G; AUPRC ranged from 0.64 to 0.74). Compared to isoform log odds ratios predicted by APARENT2, a neural network trained on APA MPRA data [22], Borzoi was consistently more accurate (Figure 5H). However, the performance gap decreased when scaling APARENT2’s odds ratio predictions by the reference isoform % from PolyADB, suggesting that context is an important determinant. We further compared to a 3^*′*^ UTR-wide ensemble of variant effect predictors, namely APARENT2 for isoform effects and Saluki [23] for half-life effects (Methods). Borzoi performs better at longer distances (dAUPRC *>* 0.050 at 2, 000 bp) with a more comparable performance closer to the PAS (dAUPRC = 0.025 at 50 bp) (Figure 5I). At closer distances, the average rank of the variant effect predictions taken over all models (Borzoi, APARENT2 and Saluki) outperforms either model’s individual performance.

### Inference of splicing events from RNA coverage

Splicing is a co- and post-transcriptional process by which the spliceosome identifies well-defined motifs near exon junctions and excises the intronic region [83, 84]. Splicing often produces alternative isoforms that may be cell- or tissue-specific [85]. Qualitatively, Borzoi predicts patterns of RNA-seq coverage in exons relative to introns well, indicating that the model has learned how sequence determines RNA splicing. As an example, Figure 6A (and Supp Figure S6A) shows predicted and measured coverage centered on an exon in the SRSF11 gene. Evaluated on the top 20% of test set genes with highest variance in coverage across the span of exons and introns, the average Pearson R between predicted and measured RNA coverage was 0.91 across all genes and samples. Researchers have previously trained deep neural networks to predict annotated splice sites as a function of the surrounding sequence [15, 16]. These methods are highly effective at classifying splice sites from non-splice sites and leave little room for improvement.

**Figure 6:**
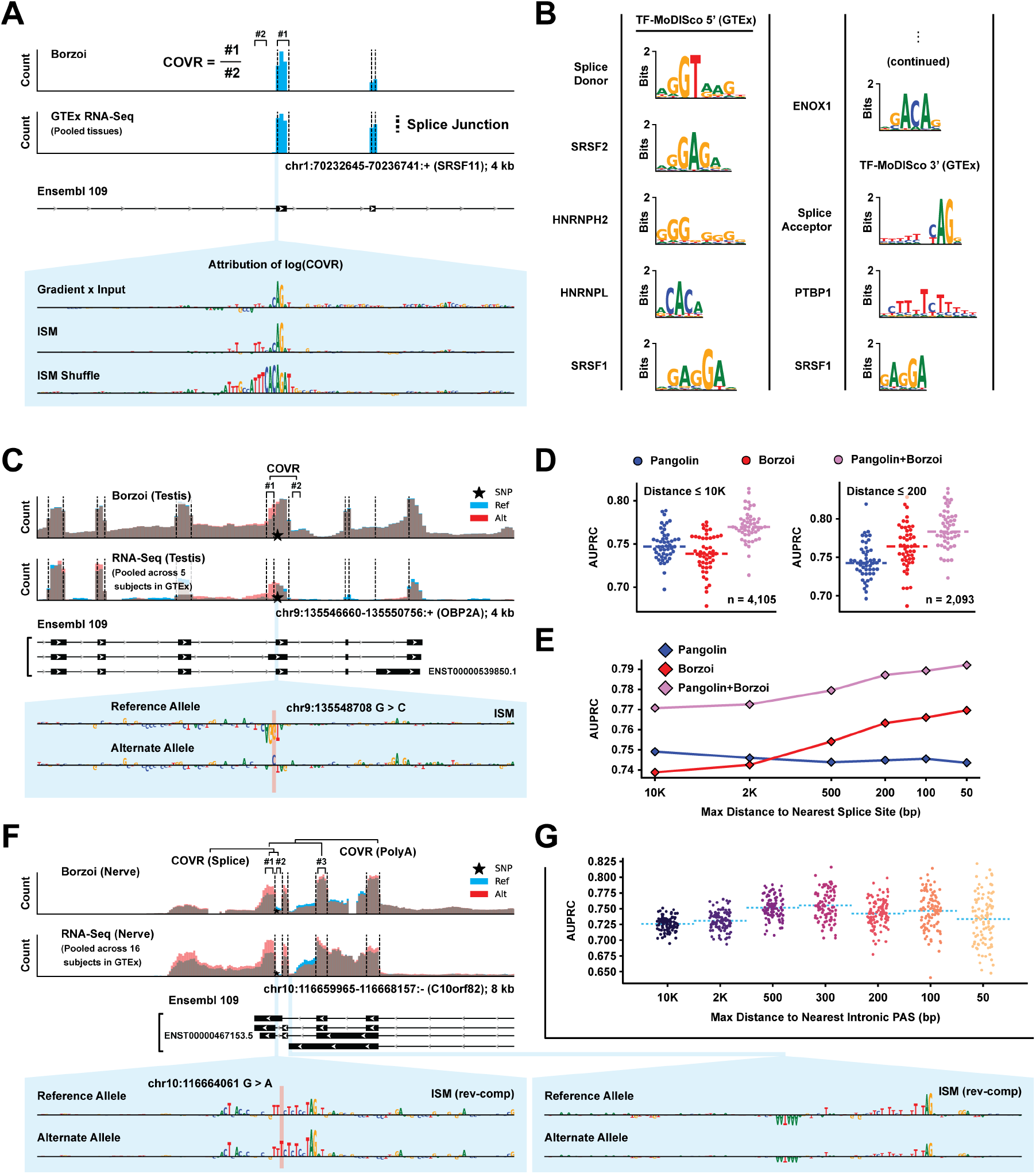
Classifying Splicing and Intronic Polyadenylation QTLs from RNA-seq coverage predictions. **(A)** Predicted and measured RNA-seq coverage across an exon in the SRSF11 gene (GTEx pooled-tissue). Calculation of exon-to-intron coverage statistics (COVR) is illustrated in the figure. Attribution scores (gradient saliency, ISM and ISM Shuffle) are shown below. **(B)** PWMs of putative splicing regulators, obtained by running MoDISco on pooled GTEx coverage ratio gradients. **(C)** Predicted RNA-seq coverage (GTEx tissue ‘Testis’) for variant rs55695858, along with measured coverage in 5 individuals with the reference allele and 5 heterozygous individuals (Testis samples). Attribution scores are shown below. **(D)** Comparison between the variant effect predictions of Borzoi, Pangolin, and an ensemble of both models at the task of classifying fine-mapped splicing QTLs from GTEx, at different distance thresholds from an annotated splice junction. **(E)** Average AUPRC for Pangolin, Borzoi, and their ensemble as a function of decreasing distance threshold to the nearest slice junction. **(F)** Predicted RNA-seq coverage (GTEx tissue ‘Nerve’) for variant rs3830026 and measured coverage in 16 individuals with the reference allele and 16 individuals who are hetero- or homozygous for the alternative allele. Bottom: Attribution scores of the exon-to-intron coverage ratio (COVR Splice) and the exon-to-exon coverage ratio (COVR PolyA). **(G)** Average AUPRC for Borzoi’s coverage ratio predictions to classify fine-mapped intronic paQTLs (tissue-pooled). Each dot represents a permutation test (Methods).

Borzoi trains indirectly on splice sites, which are present in the quantitative RNA-seq coverage labels only as 32 bp bins that transition from low intron to high exon coverage, rather than explicit nucleotide-resolution labels. Nevertheless, the predicted exon-to-intron coverage ratio surrounding a potential splice site discriminates between annotated sites and negatives with accuracies on par with the state-of-the-art model Pangolin [16] (Supp Figure S6B). When running MoDISco on gradients from tissue-pooled exon-to-intron coverage ratios for genes from the Gasperini set [53], we found known splice-regulatory sequences as top-ranking motif clusters (Figure 6B). Finally, we explored tissue-specific alternative splicing, but sequencing bias and other confounders made it challenging to definitively call such events from the predictions. Borzoi predictions tended to hedge similarly across tissues rather than capture switch-like examples of tissue-specific alternative splicing (Supp Figure S6C-D). We return to this observation in the Discussion.

### Functional splicing variant effect prediction

To benchmark variant effect scores on splicing, we curated fine-mapped splicing QTLs from the eQTL Catalog and constructed expression- and splice distance-matched negatives [82]. Variant effect scores were calculated from the predictions as the maximum absolute difference in relative coverage across bins within the gene span. While this score does not perfectly isolate the effects of splicing from e.g. transcription, it becomes a powerful statistic when explicitly distinguishing between fine-mapped sQTLs and negatives with no functional impact. RNA coverage predictions for an example sQTL (rs55695858) is shown in Figure 6C (and Supp Figure S7A), along with measured coverage among 5 individuals from GTEx with or without the alternative allele. The variant weakens an alternative 3^*′*^ splice site, which upregulates extension of the corresponding exon. Borzoi correctly predicted this change, highlighting the model’s unique ability to unravel the full transcriptomic consequences of a variant.

When comparing Borzoi to the Pangolin model [16] at the task of classifying the causal fine-mapped sQTLs from matched negatives (n = 4, 105), Pangolin has a slight advantage at large distances from annotated splice sites (Figure 6D; dAUPRC = 0.01 at distances ≤10, 000 bp). Most of these far-away SNPs are de novo splice-gain mutations. In contrast, Borzoi has an advantage at distances closer to the junction (Figure 6E; dAUPRC = 0.02 at distances ≤200 bp). Importantly, the average rank prediction of both models is superior to either model alone (dAUPRC *>* 0.02). Supp Figure S7B includes a tissue-pooled benchmark with similar results. More examples are shown in Supp Figure S7C-D.

### Intronic polyadenylation variant effect prediction

Candidate polyadenylation sites frequently occur in introns, resulting in competition between the PAS and the enveloping splice junctions [74, 86]. If the splice sites are used and the intron is excised, the excised PAS can no longer be cleaved. If instead the PAS is cleaved by the 3^*′*^-end processing complex, it results in a truncated transcript with the poly(A)-tail attaching upstream of the splice acceptor. Curious as to whether Borzoi has learned this competition between distinct regulatory functions, we filtered the paQTLs from the eQTL catalog for SNPs that were closer to intronic pA sites than 3^*′*^ UTR sites and constructed a new set of expression-controlled negatives matched for intronic pA distance.

Figure 6F highlights a fine-mapped causal intronic paQTL exhibiting competition between a set of splice sites and an intronic PAS. A G*>*A change creates a stronger polypyrimidine tract, increasing splicing efficiency and the rate at which the intron is excised. Intriguingly, RNA coverage increases non-uniformly across the transcript, with a larger increase in density for downstream exons and for the exon immediately upstream of the SNP. Attribution scores of the coverage ratio between the set of exons increasing in coverage and those with no discernible increase in coverage identify a salient intronic PAS. As splicing efficiency increases, this PAS is excised more often (hence its negative attribution), resulting in fewer truncated transcripts and increased downstream coverage. These interactions across regulatory functions complicate variant interpretation, highlighting the need for an accurate joint model of post-transcriptional regulation. Borzoi is performant at the task of classifying fine-mapped causal intronic paQTLs from negatives with an average AUPRC of 0.725 (Figure 6G). Supp Figure S7E-F highlights two additional intronic paQTLs where alterations to either the splice- or polyadenylation code result in asymmetric changes to RNA coverage across the gene body.

## Discussion

In this paper, we propose a new sequence-based machine learning model, Borzoi, that moves toward unifiying transcriptional and post-transcriptional regulation. By learning to predict sequencing coverage from a vast set of RNA-seq experiments, Borzoi enables variant scoring and interpretation through multiple layers of regulation, including transcriptional activation, splicing, and polyadenylation. Borzoi demonstrates competitive performance to state-of-the-art models in classifying fine-mapped QTLs. Intriguingly, we found that combining the predictions of Borzoi with those of other models trained on orthogonal data boosts performance. Furthermore, Borzoi has learned rules of competitive interaction for some of the regulatory functions, as exemplified by our study of intronic polyadenylation QTLs. By applying sequence attribution methods to statistics derived from the predicted coverage tracks, we show that Borzoi provides both tissue-specific interpretations of enhancers driving RNA expression in normal tissues as well as post-transcriptional variant interpretations within the mRNA transcript. By directly modeling RNA-seq coverage, Borzoi provides multi-faceted attributions based on the specific coverage ratio statistic used, exposing a dynamic “differentiable RNA-seq” interface to the user.

Challenges to modeling RNA-seq coverage data remain, and Borzoi is far from perfectly capturing the full spectra of regulation. For example, we noticed that most splicing QTLs with measured tissue-specific differences were not captured well by the model, which rather tended to predict the average RNA-seq shape. Furthermore, when we compared sequence attributions of Borzoi to those of Saluki [23] (or to MPRA-style measurements of transcript stability [78]), we did not see sequence elements related to mRNA half-life. Disentangling these layers of regulation is particularly difficult in the presence of sequencing bias. For example, reads aligning with greater density at the 3’-end of transcripts [87, 88] and other confounders (e.g. GC bias) caused false positives as we attempted to classify alternatively used splice sites based on predicted coverage. We also identified 3’ paQTLs occurring in gene promoters that were predicted and measured to differentially decrease coverage across the downstream part of the gene body. However, we had trouble concluding whether these were true 3’ isoform switches or whether the model had learned to recapitulate sequencing bias. Similarly, we found fine-mapped sQTLs that were also eQTLs, where it was difficult to disentangle true differential exon inclusion induced by the variant from effects of coverage bias observed as total expression levels decreased.

In future work, we envision several directions for improvement. RNA-seq has been adopted into a diverse set of assays to focus on specific aspects of post-transcriptional regulation—e.g. CLIP-seq to measure RBP binding [89, 90], ribosomal profiling to measure translation [91, 92], and various forms to measure RNA methylation and half-lives [93, 94]. We hypothesize that adding such training data will generally improve learning the sequence basis of these regulatory processes, but also positively influence challenging cell type specific predictions like alternative splicing. Similarly, we anticipate training on experiments in which regulatory proteins have been perturbed will improve model performance generally and enable causal inference tying particular regulators to the sequence patterns mediating their functions [95, 96]. Data quantity is a critical factor in successful machine learning and we believe that adding RNA-seq, as well as other biochemical readouts, from more mammals is a viable path to increasing training data and model quality [97]. Relatedly, training on individual human genomes with matched RNA-seq data from population sequencing efforts like GTEx [33] may help calibrate variant effect predictions and enable accurate gene expression prediction across individuals in a population [67, 68]. Finally, we are eager to incorporate new efficient attention modules to boost the receptive field to megabase scale and predict at finer resolution [98].

In summary, we developed a new neural network model for predicting RNA coverage from sequence and demonstrated its performance on numerous variant interpretation tasks. Direct modeling of RNA-seq opens the door to study a wide range of experimental assays, increasing our ability to understand the impact of genetic variation on transcription, splicing, polyadenylation and other regulatory phenomena.

## Methods

### Data

The training data for this analysis consisted of a large set of human and mouse RNA-seq experiments. However, to help the model identify distal regulatory elements away from RNA transcripts, we seeded the training data with the experimental assays studied by the Enformer and Basenji models [8, 9, 13]. This includes a curated set of human and mouse CAGE assays from the FANTOM5 consortium [99, 100] and DNase and ChIP-seq from ENCODE [31], which has absorbed the Epigenomics Roadmap [38]. We processed the data slightly differently relative to prior analyses. First, we aggregated the aligned read counts here at 32 bp resolution. Second, we split the CAGE aligned reads by strand, requiring that the model predict both the forward and anti-sense coverage.

We collected 867 human and 278 mouse RNA-seq coverage tracks from ENCODE. The tracks available for download represent normalized coverage from the STAR alignment program of uniquely mapping reads [101]. Due to the relatively large dynamic range of RNA-seq, we normalized each coverage track by exponentiating its bin values by 3*/*4. If bin values were still larger than 384 after exponentiation, we applied an additional square root transform to the residual value. This set of transformations is referred to as ‘Squashed scale’ in the main text. Most experiments used a protocol to enable stranded analysis, creating a forward and anti-sense coverage track.

We supplemented these data with 89 tracks from GTEx whole-tissue samples [33], uniformly processed by the recount3 project [34]. recount3 clustered the 49 GTEx tissues into 30 meta-tissues, combining highly related physiological regions (such as regions of the brain). For each meta-tissue, we chose a subset of samples to include as training data by performing k-means clustering on the gene expression profiles of all samples with *k* = 3 (although several meta-tissues collapsed to *k* = 2). For each cluster, we chose to include the sample with minimum average distance to all cluster members. These data were processed without consideration of strand information in recount3, which means the GTEx training tracks are non-stranded while most other RNA-seq tracks are stranded. For these tracks, we scaled the aligned fragment counts by the inverse of their average length in order to weight each fragment as a single event, in addition to the exponentiation transform described above.

In contrast to previous efforts in which a single train/validation/test split is chosen, we fragmented the human and mouse chromosomes and divided these fragments into eight roughly evenly sized folds, pairing orthologous regions into the same fold. We trained four models, each with distinct held out test and validation folds. The four models allowed us to evaluate metric variance and construct an ensemble predictor that generally performs better than any individual model.

### Model

The model was based on the Enformer network architecture but introduced a number of simplifications and enhancements to optimize for RNA-seq prediction [13]. Supp Figure S8 shows the full architecture. Enformer comprises two main stages. First, repeated application of a convolution block that achieves a 2-fold reduction of the sequence length extracts local sequence patterns until each position in the sequence represents 128 bp. Second, repeated application of a self-attention (or transformer) block enables long-range interaction and exchange between every pair of sequence positions [27, 28]. Enformer accepts a 196 kb input sequence and predicts coverage data aggregated at 128 bp resolution.

RNA-seq is a base-resolution readout of transcribed RNAs. We believed it was important to both increase the sequence length and decrease the prediction resolution to model RNA-seq well. Mammalian genes regularly exceed a full span *>* 100 kb, and if the 5^*′*^- or 3^*′*^-end of a gene extends outside of the training sequence window (such that its promoter and other regulatory signals are not captured in the receptive field of the network), it will likely obstruct learning. Conversely, mammalian exons regularly cover fewer than 128 bp, and modeling the coverage patterns around these exons at such a coarse resolution can obstruct splice site learning. However, computational limitations make these joint objectives challenging. Thus, we aimed for a compromise of 524 kb input sequences, predicting at 32 bp resolution.

Halting the convolution- and pooling blocks in the vanilla Enformer architecture at 32 bp would mean that the self-attention blocks processed 16, 384-length sequences. These blocks require quadratic memory complexity, which exceeds the capability of contemporary GPU/TPU hardware without complicated optimizations. Thus, we chose to remain at 128 bp resolution for the self-attention blocks. To predict at 32 bp resolution, we instead make use of U-net upsampling techniques from the image segmentation and object detection literature [29, 30], which solve an analogous problem of determining image-level content and communicating it back down to pixel resolution annotations. Briefly, the output embeddings predicted by the self-attention blocks at 128 bp resolution are upsampled 2x by duplicating the embedding vector at each position. We then apply point-wise convolutions in order to match the number of channels to those of the original convolution tower output (preceding the self-attention blocks) at 64 bp resolution. Finally, we add the upsampled feature map from the self-attention blocks and the intermediate feature map from the convolution tower and apply a separable convolution with width 3. This workflow is repeated once more using the intermediate feature map with 32 bp resolution from the convolution tower.

Because this architecture is still very computationally expensive, we simplified several Enformer components. First, we used max pooling instead of attention pooling, which requires an additional convolution but generally only minimally boosts performance. Second, we apply only a single width 5 convolution in each block of the initial convolution tower, forgoing the second convolution added in with a residual connection used by Enformer. Third, we reduced the number of self-attention blocks from 11 to 8 to reduce memory usage. Fourth, we used only central mask relative position embeddings since additional distance functions minimally affected performance.

### Training

Similar to previous work, we trained the model in a multi-task setting to predict coverage for all assays from one species, with a species-specific head attached to the shared model trunk. During training, we alternate human and mouse training batches by dynamically swapping in the corresponding species-specific head. In order to avoid less accurate predictions on the sequence boundaries (due to asymmetric visibility), we cropped from each side to focus the loss computation on the center 196, 608 bp. We used a Poisson loss function, but decomposed the loss analogous to BPnet to separate magnitude and shape terms [7]. Independent Poisson distributions at each sequence position is mathematically equivalent to a single Poisson distribution representing their sum, followed by allocating the counts to sequence positions using a Multinomial distribution. Thus, we apply a Poisson loss on the sum of the observed and predicted coverage and a Multinomial loss on the normalized observed and predicted coverage across the sequence length. This decomposition allows us to weight the Multinomial shape loss by a greater amount (5x), which we found boosts performance.

Using TensorFlow, backpropagation of this model on a 524 kb sequence maxes out the 40 GB of RAM of a standard Nvidia A100 GPU. We trained each model replicate using the Adam optimizer with batch size 2, split across two GPUs for *∼* 25 days and stopped training when the validation set accuracy plateaued.

### Enformer comparison

Our research objective was to extend this modeling framework to new data, i.e. RNA-seq, and not to exceed Enformer performance on the set of overlapping tracks, which includes CAGE, DNase, ATAC and ChIP assays. Several modeling decisions make this comparison imperfect. First, working with larger sequences divided into multiple folds required reprocessing the genome so that no Borzoi fold exactly matches the Enformer test set. Second, we aggregated the data at 32 bp resolution, while Enformer works with 128 bp. Third, we split the aligned reads from the CAGE datasets by strand. Nevertheless, we examined test accuracies for Borzoi versus Enformer on these overlapping datasets and found them to be broadly similar despite the modifications described above (Supp Figure S1A-D).

### Input sequence attribution

To visualize and quantify salient features in the input sequence (such as transcription factor or RNA binding protein motifs), we apply a number of different attribution methods, each of which have their own strengths and limitations. In summary, we either use methods based on (1) gradient saliency, which are computationally efficient for single outputs but tend to be noisier due to moving off the one-hot coding simplex, or (2) in-silico mutagenesis, which often give better-calibrated attributions for all outputs, but is too computationally expensive to run on long sequences. The shared goal of these methods is to estimate the contribution of each nucleotide in the input with respect to scalar statistics derived from the predicted coverage tracks, resulting in a matrix ***s*** ∈ ℝ^524,288*×*4^ of saliency scores for each coverage track. In this study, we focus solely on interpreting Borzoi’s RNA-seq tracks. Furthermore, by computing distinct summary statistics from the predicted RNA coverage tracks, we dynamically isolate distinct regulatory mechanisms in the attribution scores, namely transcription, polyadenylation and splicing.

As preliminaries, let ℳ be the Borzoi model, ***x* ∈ {**0, 1} ^524,288*×*4^ be the one-hot coded input sequence, ***y*** = ℳ(***x***) ∈ (0, +∞]^16,384*×*7,611^ be the (human) coverage prediction and let 𝒯 = {*t*_0_, …, *t*_*T*_} be the set of *T* indices of the coverage tracks in ***y*** that we want to average over (e.g. to combine all blood-specific tracks) and compute the attribution scores with respect to. Note: Borzoi’s raw prediction ***y*** is based on training data that had been subjected to various transforms intended to stabilize training (exponentiating by 3*/*4, additional exponentiation of residuals above a target value, and re-scaling). Here we assume that we have applied the inverse transforms to ***y*** such that the tensor can be reasonably assumed to reflect counts. (Also note that these transforms are differentiable, which means gradient saliency can be propagated through the inverse operations.)

### Summary statistics

1. **Log-sum of exon coverage (expression attribution):** The summary statistic *u* ∈ ℝ is computed by aggregating the set of 32 bp bins *ℬ* = *{b*_0_, …, *b*_*B*_*}* in ***y*** overlapping the exons of the gene of interest:

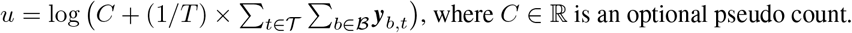
2. **Log-ratio of PAS coverage (polyadenylation attribution):** The statistic *u* ∈ ℝ is computed by summing coverage in 5 adjacent bins immediately upstream of the bin *b*_prox_, which overlaps the PAS of interest, and dividing by the coverage of a matched set of bins upstream of bin *b*_dist_ where a competing PAS is located (or alternatively immediately downstream of *b*_prox_ if the gene of interest is not subject to alternative polyadenylation):

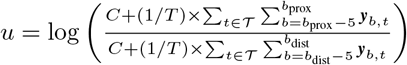 Note: The formula above assumes that the gene is on the forward (plus) strand. Coverage must be summed from *b*_prox_ + 1 to *b*_prox_ + 5 + 1 (and from *b*_dist_ + 1 to *b*_dist_ + 5 + 1) if the gene is on the minus strand.
3. **Log-ratio of exon-to-intron coverage (splicing attribution):** The statistic *u* ∈ ℝ is computed by summing coverage in bins ℬ_exon_ = {*b*_0_, …, *b*_*E*_} overlapping the exon and dividing by the sum of coverage in a matched number of bins ℬ_intron_ = {*b*_0_, …, *b*_*I*_} overlapping the adjacent intron, or alternatively a neighboring exon (which occasionally resulted in less noisy attributions when intronic polyadenylation sites or other phenomena created non-uniform intronic coverage):

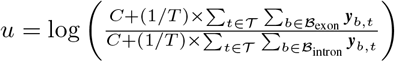

### Attribution methods

1. **Gradient x Input (‘Gradients’):** [40] Given summary statistic *u*(***x***), the attribution scores ***s* ∈** ℝ^524,288*×*4^ are computed by taking the gradient with respect to input ***x*** and subtracting the mean at each position across the four nucleotides:

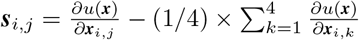 When visualizing ***s***, we extract the score at position *i* corresponding to the reference nucleotide *j* only (which is easily implemented by multiplying with ***x*** and aggregating across nucleotides):

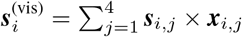
2. **Smoothgrad:** [42] We use a discretized but intuitively similar version of Smoothgrad (which originally relied on gaussian noise; here we use categorical noise). The attribution scores ***s* ∈** ℝ^524,288*×*4^ are computed as follows: Create tensor 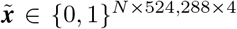 containing *N* copies of input pattern ***x*** and randomly (and independently) mutate each position in each copy with probability *σ*:

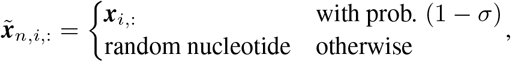

where ‘:’ refers to the entire one-hot coded dimension. Compute noisy, mean-subtracted, attributions 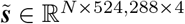 as:

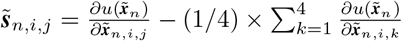 Finally average 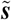 across the *N* samples and broadcast the final scores to the shape of ***s***:

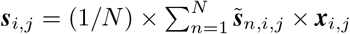
3. **In-silico Saturation Mutagenesis (‘ISM’):** Given a start- and end position *p*_start_ and *p*_end_ in ***x*** to compute ISM over, the attribution scores ***s* ∈** ℝ^524,288*×*4^ are computed as follows: Create a new tensor 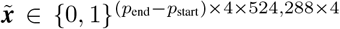 and let each matrix 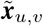 hold a mutated copy of ***x*** where the reference nucleotide at position *u* is substituted for nucleotide *v*. Then compute the ISM scores ***s*** as:

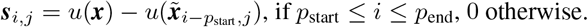 When visualizing ***s***, we average the scores across the four nucleotides:

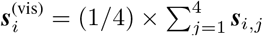
4. **Window-shuffled ISM (‘ISM Shuffle’):** Given a start- and end position *p*_start_ and *p*_end_, a window size *M* and a number of re-shuffles *N*, the attribution scores ***s*** ℝ^524,288*×*4^ are computed as follows: Create tensor 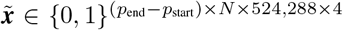 containing (*p*_end_ − *p*_start_) × *N* copies of input pattern ***x***. For each matrix 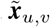 (where *v* denotes one of *N* independent samples), either dinucleotide-shuffle the local region [*u − M/*2, *u* + *M/*2 + 1] or replace the reference nucleotides in this region with uniformly random nucleotides. Dinucleotide-shuffling (with *M* = 7 and *N* = 24, or *N* = 8 for large window sizes) is performed when computing enhancer saliency while uniform random substitution (*M* = 5 and *N* = 24, or *N* = 8 for large window sizes) is used for promoters, splice sites and polyadenylation signals (where salient features are often stretches of repeating nucleotides). Then compute the attribution scores ***s*** as:

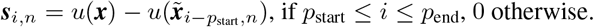 When visualizing ***s***, we average the scores across the *N* samples:

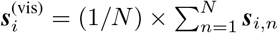

### Tissue-specific motif discovery

We visualized learned tissue-specific cis-regulatory motifs driving RNA coverage in GTEx tracks through a combination of (1) picking a large set of (measured) highly tissue-specific genes, (2) computing their gradient saliencies and normalizing out tissue-shared saliency, and (3) clustering and annotating the saliency scores using TF-MoDISco [45] and Tomtom MEME suite [46]. We first downloaded measured TPMs for GTEx v8 (GTEx_Analysis_2017-06-05_v8_- RNASeQCv1.1.9_gene_median_tpm.gct.gz). We heuristically cleaned the data by adding a small pseudo-TPM that was roughly the 1st percentile of all values (to avoid zeros), followed by clipping at a value slightly larger than the 99th percentile per tissue (to avoid extremely large numbers). Then, for each of the 5 prospective GTEx tissues ‘Whole Blood’, ‘Liver’, ‘Brain - Cortex’, ‘Muscle - Skeletal’ and ‘Esophagus - Muscularis’, we computed gene-specific log fold changes of TPM expression for the tissue of interest relative to the average TPM expression of the four other tissues. For each tissue, we sorted the TPM matrix in descending order of this metric and selected the top 1, 000 most differentially expressed genes, resulting in a total of 5, 000 genes.

We computed nucleotide-level attribution scores (input gradients) with respect to the log of aggregated exon coverage for each of the 5, 000 genes, repeating the gradient computation for each of the five GTEx tissues. Specifically, we matched each GTEx tissue to the corresponding 2-3 RNA coverage tracks obtained from recount3 and trained on (e.g. for ‘Brain - Cortex’, we compute the input gradient saliency with respect to the three GTEx ‘Brain’ super-group tracks). The gradient computation was repeated for all four model replicates, for both forward- and reverse-complemented input sequences, and averaged.

The gradient computation outlined above produces 5 separate sets of saliency scores for all 5, 000 genes (one set of scores per tissue). Next, we performed de novo motif discovery for tissue ‘x’ by slicing out the 1,000 genes originally selected to be differentially upregulated in tissue x and running TF-MoDISco on the residual gradient scores for tissue x. The residual scores were calculated by subtracting the average gradient of the four other tissues from those of tissue x, thus dampening the saliency of shared regulatory motifs and accentuating motifs specific to tissue x. Additionally, before running MoDISco we first re-weighted the gradients by computing the standard deviation at each position across the four nucleotides, applying a gaussian filter (std = 1, 280, truncate = 2) to the resulting vector of standard deviations and dividing the gradient scores by this smoothed vector. This operation results in down-weighting of regulatory regions with long contiguous stretches of large magnitude (often promoter regions) and up-weights sparser regulatory regions (transcriptional enhancers). To increase computational efficiency, we extracted the centered-on 131 kb gradient scores (as opposed to the full 524 kb) before calling MoDISco. TF-MoDISco was executed with the following parameters: ‘revcomp = true’, ‘trim_to_window_size = 24’, ‘initial_flank_to_add = 8’, ‘sliding_window_size = 18’, ‘flank_size = 8’, ‘max_seqlets_per_metacluster = 40, 000’. Other parameters were kept at their default values.

The five tissue-specific MoDISco result objects were filtered and pooled as follows: Tomtom MEME was used to match the PWMs of each MoDISco cluster to HOCOMOCO v11 [47] motifs (each PWM was trimmed by an information content threshold *>* 0.1). Only matches with E-value ≤0.1 were retained. The match with lowest p-value was chosen as the representative motif for that cluster. The five MoDISco objects were pooled by matching clusters with identical HOCOMOCO motifs and merging the seqlet coordinates, resulting in a single list of seqlet coordinates for each putative motif. A scalar tissue-specific saliency score was then computed for each seqlet by averaging the input-gated gradients overlapping its coordinates. The distributions of these seqlet-level gradient saliencies were used to assess the tissue-specificity of each motif.

Replicating the entire analysis with pseudo counts added to the predicted sum of exon coverage before applying log and computing gradients resulted in nearly identical results. Replicating the analysis without running TF-MoDISco on residual attribution scores, but rather using the raw gradients from each tissue-specific coverage track as input to TF-MoDISco, similarly produced negligible differences.

### Tissue-pooled splice motif discovery

Splice-regulatory motifs were generated by computing input gradients with respect to the splicing attribution statistic (log-ratio of exon-to-intron coverage) for one randomly chosen exon in each of the 4, 778 genes from the Gasperini et al. data [53]. The gradients were computed with respect to the average predicted coverage taken across all 89 of Borzoi’s GTEx RNA-seq tracks. The gradients were normalized across genes as follows: We first compute the standard deviation across the four nucleotides and find the maximum standard deviation across all 524, 288 positions per gene. We clip the lower end of the 4, 778 maximum deviations at the 25th percentile (to avoid up-weighting gradients with very low magnitudes) and divide each gene’s gradient by this number. Finally, to obtain 5^*′*^ splice motifs, we extracted a 192 bp window centered on the splice donor from each of the gradients. To obtain 3^*′*^ splice motifs, we extracted a 192 bp window around the splice acceptor.

TF-MoDISco was executed on the resulting 4, 778 × 192 × 4 hypothetical scores, using custom parameter settings that we empirically found worked better for degenerate RNA binding protein motifs: ‘revcomp = false’, ‘trim_to_- window_size = 8’, ‘initial_flank_to_add = 2’, ‘sliding_window_size = 6’, ‘flank_size = 2’, ‘max_seqlets_per_metacluster = 40, 000’, ‘kmer_len = 5’, ‘num_gaps = 2’, ‘num_mismatches = 1’.

### Tissue-pooled polyadenylation motif discovery

Salient motifs related to polyadenylation signals (PASs) were obtained similar to the procedure for splice-regulatory motif discovery. We computed tissue-pooled gradients with respect to the polyadenylation statistic (log-ratio of PAS coverage) for the distal-most PAS of each gene from the Gasperini set [53]. The gradients were normalized by the (clipped) maximum standard deviation per gene. Finally, a 192 bp window centered on the mode of saliency in the 3^*′*^ UTR of each gene was used to extract short gradient slices. These gradient slices were used as hypothetical scores for TF-MoDISco, which was executed using the same custom parameters as was used for splice motif discovery.

### Attention matrix visualization

We visualized higher-order structures and long-range interactions learned by Borzoi directly through the attention score matrices of the self-attention layers. Examples of such higher-order structures include intronic and exonic regions, untranslated regions (UTRs), promoters and gene spans. Long-range interactions describe relationships or dependencies between these structures learned by Borzoi, which would be observed as off-diagonal intensities in the attention matrix. Such examples include phenomena where an intron attends to its nearest exon junction, where a 3^*′*^ UTR attends to its polyadenylation signals, or where gene spans attend to promoters and transcriptional enhancers. After exploring the predicted attention maps for several different loci, we noticed that higher-order structures matching GENCODE annotations [36] were generally found in the later self-attention layers. However, to mitigate capturing potential assay- or experiment-specific biases and focus on general knowledge, we decided to not use the two final attention layers and instead used the two penultimate self-attention layers for all analyses. We further noted that different attention heads tended to capture mostly the same trends, leading us to analyze the mean attention of all 8 heads.

Let 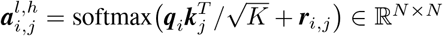 be the attention matrix for head *h* of layer *l*, where ***q***_*i*_ is the *i*:th query vector, ***k***_*j*_ is the *j*:th key vector, ***r***_*i,j*_ is the positional encoding and *K* is the key/query size. We obtain t tention matrix to be visualized as an unweighted average of all heads of the two penultimate layers: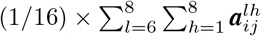. When zooming in on smaller sections of the attention matrix, we apply a small Gaussian filter to smooth out high-frequency noise (*σ* = 0.5, truncate = 2.0). We further average the attention matrix over 4 independent model replicates and reverse-complemented input sequences. Promoters generally had higher-magnitude attention values than exons, leading us to clip individual entries in the average attention matrix at 0.005 (each row of 4, 096 entries sum to 1.0).

### Fine-mapped eQTL classification and regression tasks

Expression quantitative trait loci (eQTL) studies deliver valuable data for evaluating whether Borzoi identifies the correct nucleotides driving expression and their sensitivity to specific alternative alleles. We studied GTEx v8 eQTL results from 49 tissues of varying sample sizes. We made use of summary statistics and fine-mapping results generated with SuSiE in the study by Wang et al. (2021) [1]. Only fine-mapped causal eQTLs with a posterior causal probability (PP) ≥0.9 were kept as positives. We focused all analyses on single nucleotide variants only because insertions and deletions (indels) introduce technical variance due to shifted prediction boundaries, which we aspire to alleviate in future work. In order to visualize the measured RNA-seq coverage tracks in individuals with or without the minor allele(s) of interest, we also made use of WGS genotyping data of GTEx subjects obtained through dbGAP (‘http://www.ncbi.nlm.nih.gov/gap’).

Inspired by the EMS score construction from Wang et al., who demonstrated that functional eQTL classification probabilities enable improved fine-mapping, we evaluated Borzoi and other models at the task of discriminating fine-mapped causal eQTLs from a negative set chosen to control for TSS distance. To compare against models with multiple generic outputs, we construct a feature vector based on the model predictions for each variant, and train a random forest classifier with the eQTL causal/non-causal labels. We considered a ‘SUM’ score and an ‘L2’ score to define these SNP features. For both score types, we start by centering the 524 kb input window on the SNP of interest and predict coverage ***y***^(ref)^ = *ℳ*(***x***^(ref)^), ***y***^(alt)^ = ℳ(***x***^(alt)^) ∈ ℝ^16,384*×*7,611^ for the reference and variant patterns respectively. When computing the SUM score vector ***u***(***x***^(ref)^, ***x***^(alt)^) ∈ ℝ^7,611^ for the 7, 611 distinct Borzoi tracks, we aggregate the difference between coverage predictions ***y***^(ref)^ and ***y***^(alt)^ across the length axis independently per track:

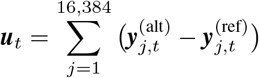

For the L2 score vector, we compute the L2 norm of the difference between predictions ***y***^(ref)^ and ***y***^(alt)^ across the length axis independently for each track. Before applying the L2 norm, we first log-transform the coverage track bins in order to focus on fold-rather than absolute change. The final metric is calculated as:

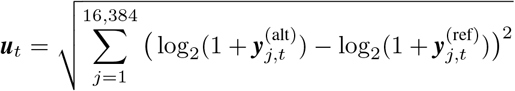

The L2 score extracts more information and achieves greater performance on this task for Borzoi. All previous Enformer work uses the SUM score, but we observed here that it also benefits from L2, though less than Borzoi.

For the second task, we evaluated models on their ability to predict eQTL effect sizes, which is a critical component of a system tasked with predicting gene expression values across a population of individuals. Because the Borzoi and Enformer models make use of gene annotation differently to map predictions to genes, we chose to perform a gene-agnostic analysis for a less biased comparison. Thus, we filtered the variant set for only those with a consistent sign of the estimated eQTL effect sizes across genes and chose the effect size with maximum absolute value as the representative effect size for that particular fine-mapped SNP. For a subset of GTEx tissues, we were able to select an appropriately matched CAGE experiment from Enformer’s outputs and computed the SUM score. For Borzoi, we selected the matching GTEx tissue RNA-seq output and computed a ‘logSUM’ score, in which we transformed the bin predictions ***y*** by log_2_(***y*** + 1) before taking a sum over the length axis.

For the third task, we evaluated Borzoi’s ability to identify the gene(s) affected by an eQTL from the set of local genes, which is intended to estimate how accurately the model can prioritize the correct gene at more general GWAS loci. We downloaded fine-mapped eQTL credible sets and their associated eGenes for 49 GTEx tissues from the eQTL catalog release 5 [81, 82]. The credible set files were downloaded from:

‘ftp://ftp.ebi.ac.uk/pub/databases/spot/eQTL/credible_sets/XYZ.purity_filtered.txt.gz’

While these file paths have since been updated, historical versions can be found here:

‘https://github.com/eQTL-Catalogue/eQTL-Catalogue-resources/blob/00ea8a7abca895f26c3aee74ece1307dc5054ace/tabix/tabix_ftp_paths.tsv’

For each variant within a credible set, we predicted a gene-specific L2 score, which considers only sequence positions overlapping the genes’ exons, for all genes within a 360, 448 bp sequence window centered on the variant. For each credible set, we computed a single score for each surrounding gene by averaging the gene’s score across variants weighted by their posterior causal probabilities. For each GTEx tissue, we computed a variant’s L2 score using model predictions for the matched GTEx RNA-seq tracks. We analyzed only credible sets associated with protein-coding genes. Due to the indel challenge described above, we further removed credible sets in which a fine-mapped variant (PP *>* 0.1) is an indel. We predicted a credible set’s target gene as the gene with the highest aggregate PP-weighted L2 score for that credible set. As a baseline, we predicted a credible set’s target gene as the nearest gene. We define “nearest gene” as the gene with the maximum PP-weighted inverse distance from the credible set. Maximizing the PP-weighted inverse distance outperforms the previously described approach of minimizing the PP-weighted distance [102]. Notably, a single distal credible set variant can inflate the minimum average distance statistic resulting in an incorrect eGene prediction, whereas maximizing the inverse distance does not suffer from this problem.

### Fine-mapped paQTL classification task

We benchmarked Borzoi’s ability to predict genetic variants that alter the relative abundance of mRNA 3^*′*^ isoforms using fine-mapped 3^*′*^ QTLs (referred to in this paper as polyadenylation QTLs, or paQTLs) obtained from the eQTL Catalog via txrevise processing [81, 82]. The file paths to the fine-mapping results were obtained from:

‘https://github.com/eQTL-Catalogue/eQTL-Catalogue-resources/blob/master/tabix/tabix_ftp_paths.tsv’

Table rows were filtered by study = ‘GTEx’ and quant_method = ‘txrev’. The resulting list of sumstat files (e.g. ‘XYZ.all.tsv.gz’) were changed to fine-map files (‘XYZ.purity_filtered.txt.gz’) and downloaded from:

‘ftp://ftp.ebi.ac.uk/pub/databases/spot/eQTL/credible_sets/XYZ.purity_filtered.txt.gz’

These file paths have since changed but a historical version of the file path table can be found at:

‘https://github.com/eQTL-Catalogue/eQTL-Catalogue-resources/blob/00ea8a7abca895f26c3aee74ece1307dc5054ace/tabix/tabix_ftp_paths.tsv’

In order to build negative sets of GTEx SNPs which are not part of any txrevise credible set, we obtained rows from the file path table where quant_method = ‘ge’ and downloaded the full sumstat files from:

‘ftp://ftp.ebi.ac.uk/pub/databases/spot/eQTL/sumstats/GTEx/ge/XYZ.all.tsv.gz‘

Fine-mapped causal paQTLs for a given tissue were obtained from the corresponding fine-mapping file (‘XYZ.purity - filtered.txt.gz’) by filtering on rows where molecular traid id contained the substring ‘.downstream.’, where the SNP occurred at most 50 bp outside of a gene span (GENCODE v41), where the distance to the nearest annotated 3^*′*^ UTR PAS in PolyADB v3 [77] was at most 10, 000 bp and where PP ≥0.9. Valid negatives were obtained from the tissue’s sumstat file (‘XYZ.all.tsv.gz’) with identical gene-span and PAS distance filters as the fine-mapped paQTLs. Negative SNPs had to either be absent from all credible sets or have PP *<* 0.01 across all GTEx tissues. Finally, we selected one negative SNP for each fine-mapped causal paQTL by requiring that they have identical distances to an annotated PAS and that the negative SNP occurs in a gene with expression levels at most 1.625-fold within the expression levels of the paQTL gene (in the same GTEx tissue). This procedure resulted in 1, 058 retained unique fine-mapped causal paQTLs.

Note that due to the relatively small number of fine-mapped paQTLs, we decided to pool all tissues, rather than benchmark separately per tissue. Since many of the positives are shared between tissues (there are a total of 1, 058 unique paQTLs, each occurring in at least one tissue), we end up with ∼2.5x the amount of unique negative SNPs after merging across tissues. Hence, for the benchmark we perform 100 permutations of randomly matching one of the multiple valid negative SNPs (from different tissues) to each corresponding positive SNP, and evaluate performance on each permutation set of 1, 058 positives and 1, 058 sampled negatives.

Intronic paQTLs (and matched negatives) were extracted from the same files as above, but had to occur in intronic regions and be closer to an annotated intronic pA site than any 3^*′*^ UTR pA site. Negatives were now matched by distance to the nearest intronic PAS. A total of 567 fine-mapped causal intronic paQTLs were retained.

### Polyadenylation variant effect prediction

We compute polyadenylation-centric variant effect scores from Borzoi’s predicted RNA coverage tracks as the maximum ratio of coverage fold change between any annotated 3^*′*^ cleavage junction within the UTR of the same gene as the SNP. Specifically, we center the 524 kb input window on the SNP, predict coverage tracks ***y***^(ref)^ = ℳ(***x***^(ref)^), ***y***^(alt)^ = ℳ(***x***^(alt)^) ∈ ℝ^16,384*×*7,611^ given the reference and alternate allele sequences ***x***^(ref)^ and ***x***^(alt)^ as input, and compute the statistic ***u***(***y***^(ref)^, ***y***^(alt)^)_*t*_ for coverage track *t* as follows:

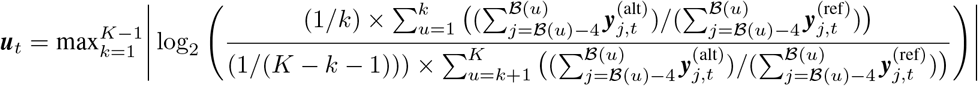

*K* in the equation above denotes the total number of polyadenylation signals within the UTR. ℬ = {*b*_0_, …, *b*_*K*_} is the ordered set of bin indices in ***y*** overlapping the *K* polyadenylation signals. The final score used in the benchmarks was the average statistic computed from all of Borzoi’s 89 GTEx coverage tracks. The score was also averaged over all four model replicates in both forward and reverse-complemented input format.

### Comparison to APARENT2 and Saluki

We compare Borzoi’s classification performance to APARENT2 in two ways: First, we score the reference and alternate PAS sequence affected by the variant using APARENT2 and simply use the absolute value of the predicted log odds ratio as the variant effect score. Second, we use the predicted odds ratio to scale the tissue-pooled reference PAS usage (as reported in PolyADB) and use the absolute value of the difference in PAS usage as the final variant effect score. The latter statistic effectively dampens the magnitude of variants which, based on APARENT2’s prediction, has a large predicted fold change, but according to measurements occur in lowly utilized PASs (due to competing PASs).

When comparing performance to an ensemble consisting of both APARENT2 and Saluki on the paQTL classification task, we follow the methodology from the APARENT2 paper [22]. Briefly, we curate the PAS sequences and corresponding mRNA isoforms of each gene (at most 30) based on annotations from PolyADB and fit a logistic regression model to predict tissue-pooled distal isoform proportions (as reported in PolyADB) given both APARENT2’s PAS scores (at most 30 scalars) and Saluki’s isoform scores (at most 30 vectors of top-4 PCA components extracted from the penultimate layer of Saluki) as input. Using this calibrated ensemble model, we predict the reference and alternate distal proportions of a gene when inducing a particular variant (which may affect multiple PAS- and isoform sequences). We estimate a final odds ratio from the predicted distal proportions and use the odds ratio to recalculate the alternate distal proportion based on the measured reference distal proportion. Finally, we subtract the alternate distal proportion from the reference proportion and use the absolute value of this difference as the final variant effect score.

### Fine-mapped sQTL classification task

Fine-mapped sQTLs and matched negatives were obtained from the eQTL Catalog [81, 82] using the same sumstat (‘XYZ.all.tsv.gz’) and fine-mapping (‘XYZ.purity_filtered.txt.gz’) files as were used for the paQTL classification task. The fine-mapped causal sQTLs were extracted by filtering on rows where molecular trait id contained the substring ‘.contained.’. These QTLs were further filtered on PP ≥0.9 and on a maximum distance ≤10, 000 bp to an annotated splice junction (GENCODE v41). A set of expression- and distance-matched negatives were constructed per tissue in an identical fashion to the paQTL task, with the exception of matching by nearest distance to splice junctions. We retained a total of 4, 105 unique fine-mapped causal sQTL SNPs.

### Splicing variant effect prediction

Purely isolating splicing impact from other mechanisms proved challenging. We focus on a simple statistic that worked well in practice, namely the maximum difference in normalized coverage across the gene span. Specifically, we center the 524 kb input window on the SNP, predict coverage tracks ***y***^(ref)^ = ℳ(***x***^(ref)^), ***y***^(alt)^ = ℳ (***x***^(alt)^) ∈ R^16,384*×*7,611^ and compute the statistic ***u***(***y***^(ref)^, ***y***^(alt)^)_*t*_ for coverage track *t* as follows:

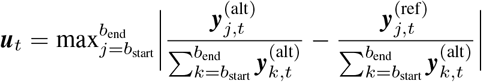

The indices *b*_start_ and *b*_end_ in the bove equation refer to the bins in ***y*** overlapping the start- and end positions of the gene span. The relatively large number of fine-mapped causal sQTLs allows for a tissue-specific benchmark comparison. To that end, for a given SNP and GTEx tissue we average the computed statistic over only the subset of predicted coverage tracks corresponding to the tissue.

### Comparison to Pangolin

We used the pre-packaged command-line utility to score sQTL SNPs with Pangolin [16]. To make comparisons easier, we modified the program to output scores with 6 rather than 2 decimals. We used the following command to score the positive and negative vcf files:

~~~
pangolin -d 2000 -m False <sqtl file>.vcf hg38.fa gencode41_basic_nort_protein.db <out_dir>
~~~

While this command allows at most a distance of 2, 000 bp from an annotated splice junction, Pangolin will also score potential de novo splice gains at the variant position, meaning that the command will produce variant effect scores for all variants (even those separated by *>* 2, 000 bp from a splice site). We parsed the command-line output and matched the gene identifier of the Pangolin output to the gene that the SNP occurs in. The final variant effect score is calculated as the sum of the absolute values of the predicted maximum increase and decrease.

### Splice site identification task

Identifying splice sites in DNA sequences has formed the basis for a successful approach to interpret the splicing code and prioritize pathogenic splicing variants [15, 16]. To evaluate Borzoi’s ability to identify splice sites, we constructed an analogous classification task and compared to Pangolin [16]. We downloaded the splicing junction counts for all GTEx samples from recount3 and selected positive examples from annotated junctions with coverage above the 50th percentile of aligned read counts. We filtered this set for those that fall in Pangolin’s test chromosomes and outside Borzoi’s third fold training regions (which had the maximal overlap with Pangolin’s test among the folds). For each positive example, we selected a matching negative site that has the same tri-nucleotide context, is between 100-2, 000 bp away, and lacks evidence for being a splice junction itself. For Borzoi, we scored each site as the predicted log coverage ratio on the exonversus intron side of the junction, averaged across samples from the corresponding GTEx tissue. For Pangolin, we scored each site with its predicted splice site probability, averaged across all tissues.

### Classifying rare and common variation from gnomAD

We sampled a set of 14, 198 singletons and 14, 198 matched common variants (allele frequency *>* 5%) from the GnomAD v3.1 database (https://gnomad.broadinstitute.org), with sampling restricted to regions overlapping ENCODE cCREs. To control for sequence mutability, we excluded variants within CpG islands and low-complexity regions. For each singleton sampled, we sampled a matched common variant with the same reference and alternate allele as the singleton. We also matched the variants’ background DNA contexts, sampling common variants that lie within the same tri-nucleotide as the singleton. Finally, we removed variants overlapping gene exons in coding sequences (GENCODE v41), focusing only on regulatory variants for our evaluation. For all sampled variants, we used their CADD raw score and CADD phred scores (v1.6) from the GnomAD v3.1 dataset. We trained ridge regression models to discriminate common variants from singletons and used 10-fold cross-validation to evaluate the models. The CADD-based model uses the CADD scores as features, whereas the Borzoi-based model uses the L2 scores across all RNA-seq tracks as features, averaged across the four training fold models. We derived a third (combined) model by averaging predicted variant ranks for the Borzoi-based and CADD-based models. For a second genome-wide second benchmark, we sampled uniformly from across the genome instead of restricting the variant sampling to ENCODE cCRES. This resulted in a variant set containing 17, 360 singletons and 17, 360 matched common variants.

### Predicting TRIP expression

We downloaded TRIP insertion coordinates and measured expression levels for 7 distinct promoters from the supplementary material of Leemans et al. (2019) [62]. The promoter sequences are listed in Table S1 and the insertion coordinates (and measurements) are listed in Data S2 of their paper. To predict the activity of TRIP reporters, we iterated over each promoter sequence and coordinate, centered the 524 kb input window on the insertion coordinate and inserted the sequence. When deriving statistics from Borzoi’s RNA-seq or CAGE predictions, we inserted the entire TRIP reporter into the genomic location (including the promoter sequence, the GFP CDS, the PAS, and the PiggyBac terminal repeat regions). In contrast, when deriving statistics from Borzoi’s DNase or histone modification tracks (e.g. H3K4me3) we only inserted the promoter, as these predictions became marginally worse when inserting the full reporter. We attribute this phenomenon to the transposable elements flanking the reporter, which Borzoi inherently does not predict well due to clipping of unmappable regions during the original training data processing.

Given the predicted coverage ***y*** = ℳ(***x***) ∈ ℝ^16,384*×T*^ for the *T* coverage tracks considered (e.g. K562 DNase tracks), we calculate a scalar prediction *u*(***x***) ∈ ℝ by averaging the coverage tracks, aggregating the signal in a local window of size *W* centered at the insertion site, and apply a log2 transform:

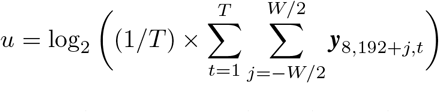

For each type of assay (e.g. DNase), we exhaustively search for the window size *W* that maximizes the spearman correlation between the resulting scalar predictions and the TRIP measurements. Note that when we perform 20-fold cross-validation, we only use the training split of the current fold of the data to search for the optimal window size.

### Gene-enhancer prioritization task

We evaluated Borzoi’s ability to link distal regulatory elements to genes by analyzing experiments in which CRISPRi was used to block the regulatory element followed by measuring gene expression. These experiments have been performed on a small set of specifically chosen genes where expression was measured by various techniques [55, 57, 58, 59, 60, 48] and a large set of all expressed genes where perturbation and expression was measured by single-cell RNA-seq [53]. These datasets were analyzed to consider whether each tested regulatory element significantly altered gene expression, defining a set of binary labels. The flow/proliferation dataset contains 117 positives out of 2, 194 tested within 262 kb of the gene’s TSS. After filtering for only genes with ≥3 elements tested, the scRNA-seq dataset contains 404 positives of 19, 104 tested within 262 kb of the gene’s TSS.

For both Enformer and Borzoi, we scored putative enhancers using input gradient analysis. For Enformer, we computed the gradient of the K562 CAGE prediction in the two 128 bp bins centered at the gene’s TSS, chosen by Enformer to have the greatest prediction for that gene. For Borzoi, we computed the gradient of the K562 RNA-seq prediction for all bins overlapping the gene’s exons in GENCODE v41. For each nucleotide, we took the absolute value of the reference nucleotide gradient. For each regulatory element, we computed a weighted average of the nucleotide scores using Gaussian weights (standard deviation 300), centered at the element’s mid point. To calibrate scores across genes with different expression levels, we normalized the scores by the mean nucleotide score across the entire region.

### Codon stability comparison

Prior work has demonstrated strong relationships between codon usage and mRNA half-life [23, 79]. We constructed a Borzoi codon statistic to compare to those previously measured. For the Gasperini scRNA-seq enhancer screen, we computed input gradients for a set of 4, 778 genes for K562 gene expression. We made use of these gradients here to quantify codon contributions to expression. For each reference codon in these genes, we used the gradients to approximate the predicted effect of changing it to all alternative codons with a single base-pair mutation. We used least squares regression to fit a coefficient for each codon on this set of possible codon mutations and effects. Finally, we compared these coefficients to codon stability coefficients computed by Forrest et al. as the Pearson correlation between codon frequency and mRNA half-life in three mammalian cell lines–HeLA, mouse ESCs, and CHO cells [79].

## Availability of data and software

The code repository for training RNA-seq deep learning models, including example code to use the model, is available under the Apache 2.0 open source license at: ‘https://github.com/calico/borzoi‘. Pre-trained Borzoi model weights are available through Github. The processed Borzoi training data (including one-hot coded sequences and coverage tracks) are available for download at: ‘gs://borzoi-paper/data/’ (Google Cloud Storage).

Gene annotations were obtained from: ‘https://www.gencodegenes.org/’ (v41). CRISPRi data was obtained from Nasser et al. (2021) and from GEO accession GSE120861 for the Gasperini et al. (2019) data. DNase, ChIP-seq, CAGE and RNA-seq data was downloaded and processed from ENCODE (‘https://www.encodeproject.org/’). Processed RNA-seq samples for GTEx individuals were downloaded from recount3 (‘https://rna.recount.bio/‘). Fine-mapped eQTLs were obtained from the supplementary material of Wang et al. (2021). Fine-mapped eQTL credible sets and other QTLs (sQTLs and paQTLs) were downloaded from the eQTL Catalog (‘https://www.ebi.ac.uk/eqtl/’). The positive (fine-mapped causal) and negative e-/s-/pa-QTL sets used in this study are available at: ‘gs://borzoi-paper/qtl/’ (Google Cloud Storage). TRIP data was downloaded from the supplementary material of Leemans et al. (2019).

The Genotype-Tissue Expression (GTEx) Project was supported by the Common Fund of the Office of the Director of the National Institutes of Health. Additional funds were provided by the NCI, NHGRI, NHLBI, NIDA, NIMH, and NINDS. The datasets used for the analyses described in this manuscript were obtained from dbGaP at http://www.ncbi.nlm.nih.gov/gap through dbGaP accession number phs000424.v9.p2.

## Author’s Contributions

Conceptualization: D.R.K.; Analysis: J.L., D.R.K., D.S., H.Y., V.A.; Writing: J.L., D.R.K., D.S., H.Y., V.A.

## Acknowledgements

We thank Anya Korsakova, Xingfan Huang, Melih Yilmaz, and Jun Xu for helpful discussions and valuable feedback.

## Funding

This work was funded by Calico Life Sciences LLC.

## Competing interests

D.R.K., J.L., D.S., and H.Y. are employees of Calico Life Sciences LLC. V.A. is an employee of Sanofi Pasteur Inc, but was involved in this work independently of Sanofi.

## Notes

https://github.com/calico/borzoi

