## Supplementary Figures 1-8 for "Predicting RNA-seq coverage from DNA sequence as a unifying model of gene regulation"

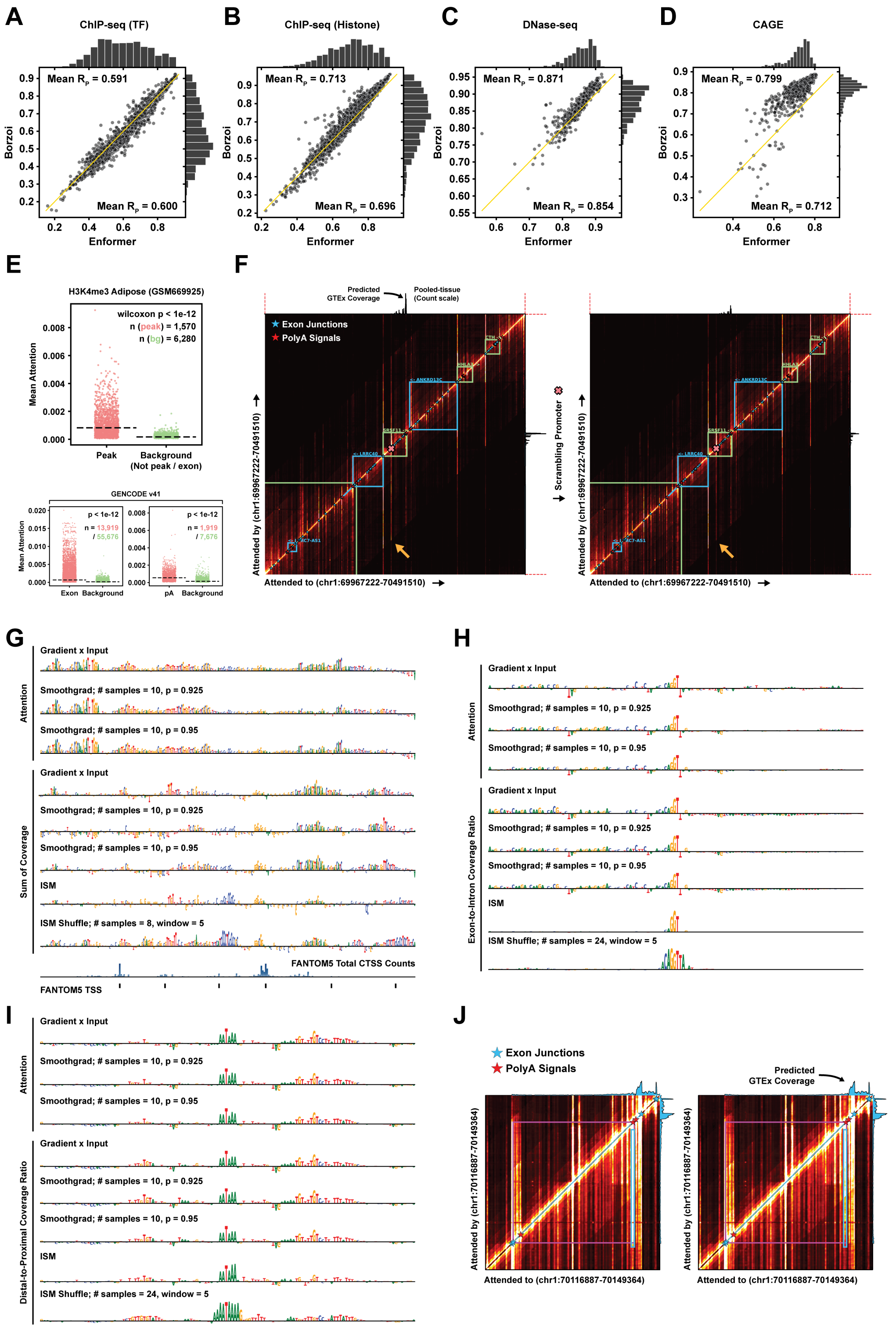

Figure S1: **Related to Figure 1.** **(A)** Performance comparison between Borzoi and Enformer on held-out genomic (human) sequences when tasked with inferring ChIP-seq (TF) coverage. Each dot in the scatter plot represents the Pearson R between predicted and observed bin-level coverage values. Note that directly comparing the performance between Borzoi and Enformer on held-out test data in this manner is not flawless, since the output bin resolution has changed, some assays (such as CAGE) have been re-processed to retain strand information, and the longer input sequence length of Borzoi changes the exact fragments contained in the train, validation and test sets compared to Enformer. **(B)** Bin-level performance comparison (Pearson R) on held-out genomic sequences when predicting ChIP-seq (histone) coverage. **(C)** Bin-level performance comparison (Pearson R) on held-out genomic sequences when predicting DNase-seq coverage. **(D)** Bin-level performance comparison (Pearson R) on held-out genomic sequences when predicting CAGE coverage. **(E)** Average attention within vertical stripes overlapping annotated H3K4me3 peaks (Adipose tissue), exonic regions, or in a 128-bp window around polyadenylation sites (as annotated in GENCODE v41). The coordinates of H3K4me3 peaks, exons and polyadenylation sites were extracted from the first 256 loci taken from the test set of the first cross-validation fold of the model. Only the first model fold were used to generate the corresponding attention weight matrices. Size-matched negative background regions were sampled from attention bins that did not overlap any peak, exon or polyadenylation site, at a rate of 4x the number of positives. P-values were calculated with two-sided wilcoxon tests. **(F)** Attention weight matrix averaged across all 8 heads of the final transformer layers, shown for example region chr1:69967222-70491510. Left attention map: Reference sequence. Right attention map: Result of dinucleotide-shuffling a 256 bp region overlapping the second TSS of the SRSF11 gene (orange arrow). Average predicted RNA-seq coverage for 89 GTEx samples is shown above each heatmap. **(G)** Attribution scores of the SRSF11 promoter (identical region as in Figure 1G), using a variety of attribution methods. The attribution scores are calculated with respect to the sum of attention overlapping the second TSS (top) or with respect to the predicted log-sum of exon coverage in GTEx 'Lung' tissue samples (bottom). Parameters specific to each attribution method are annotated above each sequence logo. **(H)** Attribution scores of one of the splice donors of the SRSF11 gene (Figure 1H). The attribution scores are calculated with respect to the log ratio of coverage across the exon relative to coverage over the intron. **(I)** Attribution scores of the distal-most PAS of the SRSF11 gene (Figure 1I). The attribution scores are calculated with respect to the log ratio of coverage immediately upstream of the distal PAS relative to coverage immediately upstream of the proximal PAS. **(J)** Average attention weight matrix displayed for the 3' UTR of the LRRC7 gene (coordinates chr1:70116887-70149364). Left: Reference sequence. Right: Result of mutating the core hexamer motif of the distal-most PAS of LRRC7.

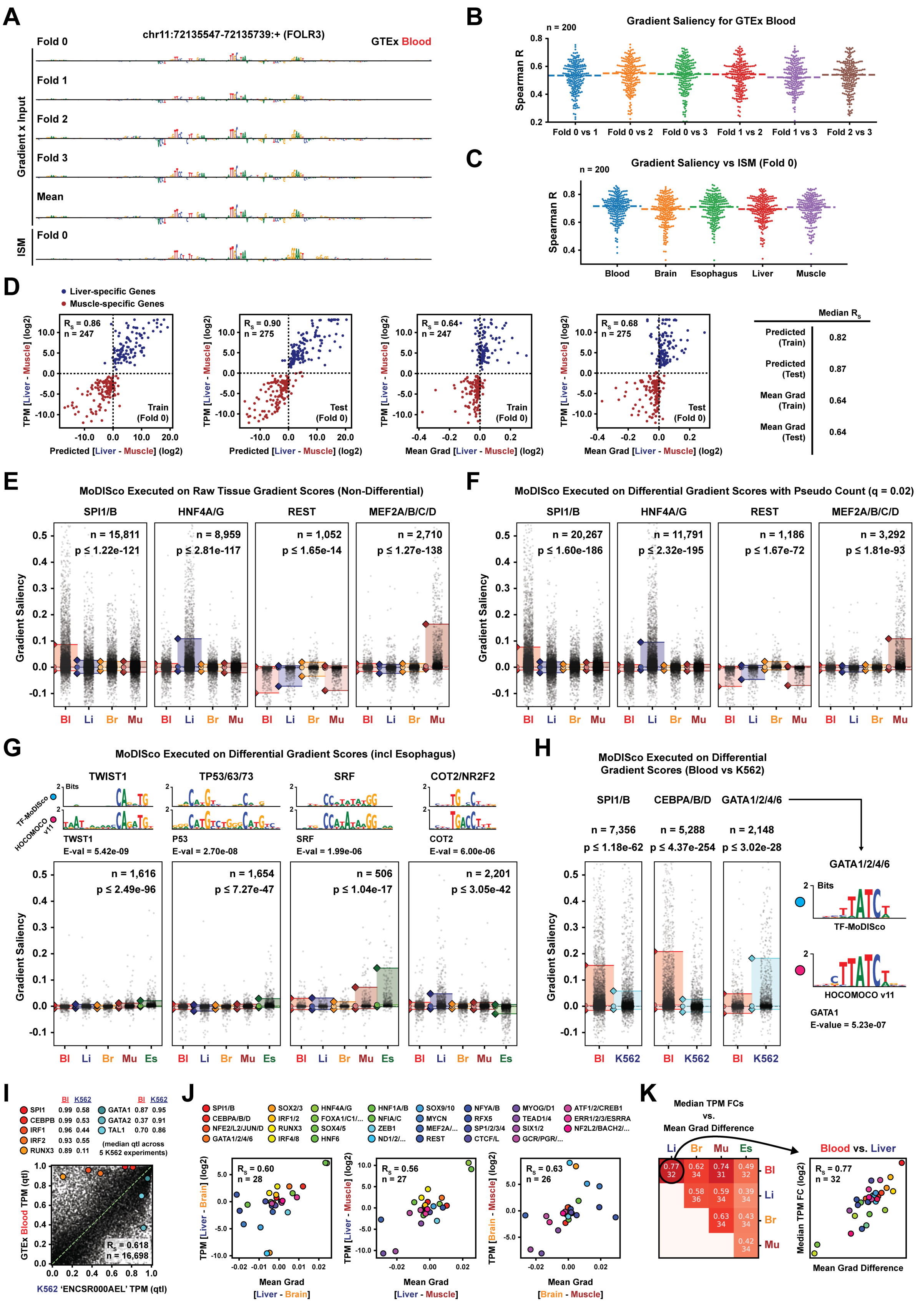
A
chr11:72135547-72135739:+ (FOLR3)
GTEx Blood
Gradient x Input
Fold 0
Fold 1
Fold 2
Fold 3
Mean
ISM
Fold 0
B
Gradient Saliency for GTEx Blood
n = 200
Spearman R
Fold 0 vs 1
Fold 0 vs 2
Fold 0 vs 3
Fold 1 vs 2
Fold 1 vs 3
Fold 2 vs 3
C
Gradient Saliency vs ISM (Fold 0)
n = 200
Spearman R
Blood
Brain
Esophagus
Liver
Muscle
D
Liver-specific Genes
Muscle-specific Genes
TPM [Liver - Muscle] (log2)
Predicted [Liver - Muscle] (log2)
Mean Grad [Liver - Muscle] (log2)
Rs = 0.86 n = 247 Train (Fold 0)
Rs = 0.90 n = 275 Test (Fold 0)
Rs = 0.64 n = 247 Train (Fold 0)
Rs = 0.68 n = 275 Test (Fold 0)
Median Rs
Predicted (Train) 0.82
Predicted (Test) 0.87
Mean Grad (Train) 0.64
Mean Grad (Test) 0.64
E
MoDISco Executed on Raw Tissue Gradient Scores (Non-Differential)
SPI1/B n = 15,811 p ≤ 1.22e-121
HNF4A/G n = 8,959 p ≤ 2.81e-117
REST n = 1,052 p ≤ 1.65e-14
MEF2A/B/C/D n = 2,710 p ≤ 1.27e-138
Gradient Saliency
Bl Li Br Mu
F
MoDISco Executed on Differential Gradient Scores with Pseudo Count (q = 0.02)
SPI1/B n = 20,267 p ≤ 1.60e-186
HNF4A/G n = 11,791 p ≤ 2.32e-195
REST n = 1,186 p ≤ 1.67e-72
MEF2A/B/C/D n = 3,292 p ≤ 1.81e-93
Gradient Saliency
Bl Li Br Mu
G
MoDISco Executed on Differential Gradient Scores (incl Esophagus)
TF-MoDISco
HOCOMOCO v11
TWIST1 TWST1 E-val = 5.42e-09
TP53/63/73 P53 E-val = 2.70e-08
SRF SRF E-val = 1.99e-06
COT2/NR2F2 COT2 E-val = 6.00e-06
n = 1,616 p ≤ 2.49e-96
n = 1,654 p ≤ 7.27e-47
n = 506 p ≤ 1.04e-17
n = 2,201 p ≤ 3.05e-42
Gradient Saliency
Bl Li Br Mu Es
H
MoDISco Executed on Differential Gradient Scores (Blood vs K562)
SPI1/B n = 7,356 p ≤ 1.18e-62
CEBPA/B/D n = 5,288 p ≤ 4.37e-254
GATA1/2/4/6 n = 2,148 p ≤ 3.02e-28
Gradient Saliency
Bl K562
GATA1/2/4/6
TF-MoDISco
HOCOMOCO v11
GATA1
E-value = 5.23e-07
I
Bl K562
SPI1 0.99 0.58
CEBPB 0.99 0.53
IRF1 0.96 0.44
IRF2 0.93 0.55
RUNX3 0.89 0.11
GATA1 0.87 0.95
GATA2 0.37 0.91
TAL1 0.70 0.86
(median qtl across 5 K562 experiments)
GTEx Blood TPM (qtl)
K562 'ENCSR000AEL' TPM (qtl)
Rs = 0.618
n = 16,698
J
SPI1/B CEBPA/B/D NFE2/L2/JUN/D GATA1/2/4/6 SOX2/3 IRF1/2 RUNX3 IRF4/8 HNF4A/G FOXA1/C1/... SOX4/5 HNF6 HNF1A/B NFIA/C ZEB1 ND1/2/... SOX9/10 MYCN MEF2A/... REST NFYA/B RFX5 SP1/2/3/4 CTCF/L MYOG/D1 TEAD1/4 SIX1/2 GCR/PGR/... ATF1/2/CREB1 ERR1/2/3/ESRRA NF2L2/BACH2/...
Rs = 0.60 n = 28
Rs = 0.56 n = 27
Rs = 0.63 n = 26
TPM [Liver - Brain] (log2)
TPM [Liver - Muscle] (log2)
TPM [Brain - Muscle] (log2)
Mean Grad [Liver - Brain]
Mean Grad [Liver - Muscle]
Mean Grad [Brain - Muscle]
K
Median TPM FCs vs. Mean Grad Difference
Li Br Mu Es
Bl 0.77 32 | 0.62 34 | 0.74 31 | 0.49 32
Li 0.58 36 | 0.59 34 | 0.39 34
Br 0.63 34 | 0.43 34
Mu 0.42 34
Blood vs. Liver
Rs = 0.77
n = 32
Median TPM FC (log2)
Mean Grad Difference

Figure S2: **Related to Figure 2.** **(A)** Gradient saliency scores calculated with respect to the log-sum of exon coverage for gene FOLR3 (GTEx 'Whole Blood' tracks), displayed in a local sequence window centered on the region with maximal value. Gradients are shown for all 4 model replicates, as well as their average. In-silico saturation mutagenesis (ISM) is shown for the first model replicate ('Fold 0') at the bottom. **(B)** Distribution of correlations (Spearman R values) between gradient saliencies of different model replicates, computed over 200 blood-specific genes (maximal fold change of 'Whole Blood' TPM relative to TPM of other tissues in GTEx). The correlations were computed in a local window of $1,024$ bp centered on the position of maximal gradient score. **(C)** Distribution of correlations (Spearman R) between gradient saliency and ISM ('Fold 0' only), computed for 200 blood-specific genes. The correlations were calculated in a local window of 192 bp centered on the position of maximal gradient score. **(D)** Left: Comparison between measured median TPM log2 fold changes in GTEx tissues 'Liver' and 'Muscle' and either (1) log2 fold changes of predicted sum of exon coverage in Borzoi's corresponding GTEx tracks, or (2) the average difference in input-gated gradient saliency within a 192-bp window centered at the mode of maximal differential saliency per gene. The predictions and gradients were computed using only the first cross-validation fold of the model. The metrics were evaluated on the subset of $5,000$ previously chosen tissue-specific genes that (by random chance) happened to belong to the test set of the first model fold (denoted 'Test' in the figures), or on a size-matched sample of trained-on loci (denoted 'Train'). Right: Median Spearman correlations of the aforementioned evaluations taken across all combinations of tissue pairs (for tissues 'Whole Blood', 'Liver', 'Brain - Cortex', 'Muscle' and 'Esophagus - Muscularis'). **(E)** A subset of motif clusters identified by MoDISco when executed on raw gradient saliencies computed from 4 GTEx RNA-seq coverage tracks ('raw' = no subtraction of the average gradient saliency of other tissue tracks; e.g. to identify blood-specific motifs, we execute MoDISco on a set of $1,000$ blood-specific genes using only the gradient saliencies derived from the 'Whole Blood' coverage predictions). The gradients were computed with respect to the log-sum of exon coverage for each target gene. Shown are the MoDISco PWMs, the best-matching HOCOMOCO v11 PWMs and the distribution of tissue-specific gradient saliencies for within-cluster seqlets. Saliency distribution p-values were computed using a two-sided Wilcoxon test between the tissue with largest absolute magnitude in gradient saliency (95th percentile) and the tissue with second largest saliency magnitude. Each score distribution is annotated with a transparent colored bar that extends to the 5th, 50th and 95th percentile of values. **(F)** The same subset of motifs identified by running MoDISco on residual gradient saliencies with pseudo count ('residual' = normalized by subtracting the average gradient saliency of other tissues from the gradients of the tissue of interest). The pseudo count was calculated as the 2nd percentile of the sum of exon coverages predicted for the subset of selected genes that we interpreted with gradient saliency ($n = 5,000$). The pseudo count was added to the predicted sum of exon coverages before taking the log (after which the gradient is computed). **(G)** Additional MoDISco motif clusters identified by interpreting Esophagus-specific genes (using predictions and gradients derived from Esophagus coverage tracks). The gradients were normalized by subtracting the average saliency of other tissues. **(H)** Motif clusters identified by MoDISco in gradients derived from Borzoi's GTEx 'Whole Blood' and K562 RNA coverage tracks, highlighting the differences in learned regulatory patterns from these two cell- and tissue states.
**(I)** Top: Median TPM quantile of a subset of blood-specific TF genes in GTEx 'Whole Blood' compared to TPM quantiles in K562 (median quantiles calculated across 5 ENCODE experiments: 'ENCSR000AEL', 'ENCSR000AEM', 'ENCSR000AEN', 'ENCSR000AEO', 'ENCSR000CPH'). Bottom: Scatter plot comparing TPM quantiles of genes in GTEx 'Whole Blood' to K562 measurements (ENCODE experiment 'ENCSR000AEL'). Blood- and K562-enriched TF genes are annotated with red and blue color respectively. **(J)** Average residual gradient saliency of MoDISco cluster seqlets for pairs of 4 distinct GTEx tissues, compared to the measured difference in log-TPM of the corresponding TF genes within the same GTEx tissues ('Whole Blood', 'Liver', 'Brain - Cortex' and 'Muscle'). The median TPM of genes belonging to the same TF subfamily (according to HOCOMOCO v11) were averaged. A subset of scatter plots for this analysis were shown in Figure 2C. **(K)** Correlation between the average residual gradient saliency of MoDISco motifs and the difference in gene TF log-TPM for all 5 GTEx tissues (including 'Esophagus - Muscularis').

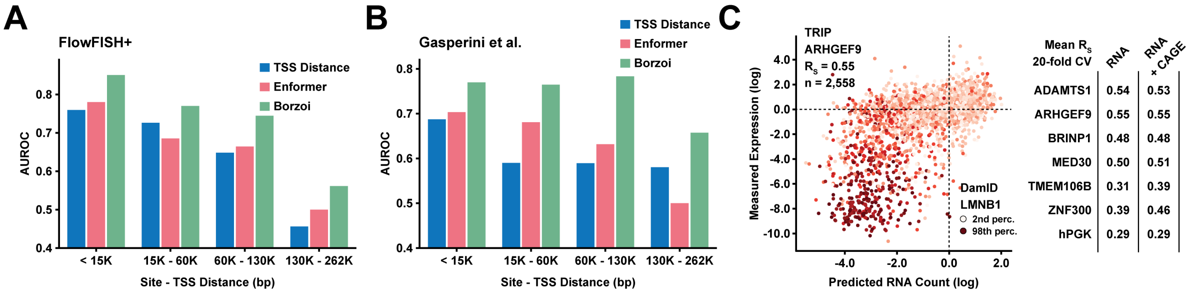


Figure S3: **Related to Figure 3. (A)** Area under the receiver operating characteristic curve (AUROC) when using a statistic computed from the Borzoi or Enformer gradient saliencies to classify whether or not a given CRE locus regulates a target gene (measurements from Fulco et al., 2016, 2019 and Klann et al., 2017, and others). The baseline performance (blue bars) corresponds to using only TSS distance when performing the classification. **(B)** AUROCs when using the Borzoi or Enformer gradient scores to classify regulating / non-regulating CRE loci in the data from Gasperini et al. (2019). **(C)** Left: Predicted vs measured expression levels of TRIP reporter constructs based on Borzoi RNA-seq coverage in K562 (Promoter: ARHGEF9). Color = DamID measurements. Right: Average Spearman R (20-fold cross validation) when predicting TRIP expression based on (log-)coverage of K562 RNA-seq tracks, or when using coverage statistics derived from both RNA-seq tracks and CAGE tracks to make the prediction (Methods).

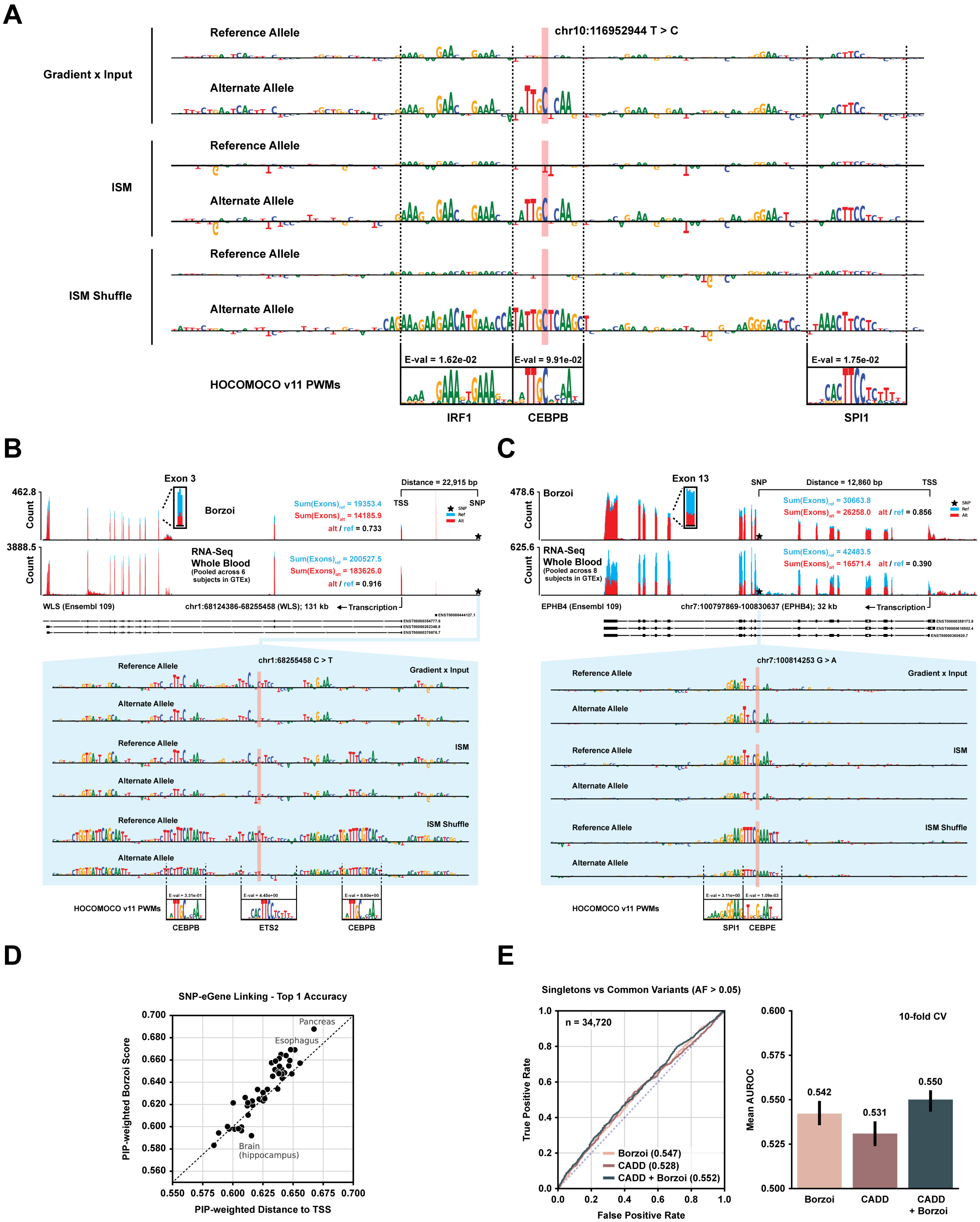

A
chr10:116952944 T > C
Gradient x Input
ISM
ISM Shuffle
Reference Allele
Alternate Allele
E-val = 1.62e-02
E-val = 9.91e-02
E-val = 1.75e-02
HOCOMOCO v11 PWMs
IRF1
CEBPB
SPI1
B
Exon 3
Distance = 22,915 bp
TSS
SNP
462.8
Count
Borzoi
Sum(Exons)ref = 19353.4
Sum(Exons)alt = 14185.9
alt / ref = 0.733
3888.5
RNA-Seq
Whole Blood
(Pooled across 6 subjects in GTEx)
Sum(Exons)ref = 200527.5
Sum(Exons)alt = 183626.0
alt / ref = 0.916
WLS (Ensembl 109)
chr1:68124386-68255458 (WLS); 131 kb
Transcription
chr1:68255458 C > T
Gradient x Input
ISM
ISM Shuffle
HOCOMOCO v11 PWMs
CEBPB
ETS2
CEBPB
C
Exon 13
SNP
Distance = 12,860 bp
TSS
478.6
Count
Borzoi
Sum(Exons)ref = 30663.8
Sum(Exons)alt = 26258.0
alt / ref = 0.856
625.6
RNA-Seq
Whole Blood
(Pooled across 8 subjects in GTEx)
Sum(Exons)ref = 42483.5
Sum(Exons)alt = 16571.4
alt / ref = 0.390
EPHB4 (Ensembl 109)
chr7:100797869-100830637 (EPHB4); 32 kb
Transcription
chr7:100814253 G > A
Gradient x Input
ISM
ISM Shuffle
HOCOMOCO v11 PWMs
SPI1
CEBPE
D
SNP-eGene Linking - Top 1 Accuracy
PIP-weighted Borzoi Score
PIP-weighted Distance to TSS
Pancreas
Esophagus
Brain (hippocampus)
0.550 0.575 0.600 0.625 0.650 0.675 0.700
0.560 0.580 0.600 0.620 0.640 0.660 0.680 0.700
E
Singletons vs Common Variants (AF > 0.05)
n = 34,720
True Positive Rate
False Positive Rate
Borzoi (0.547)
CADD (0.528)
CADD + Borzoi (0.552)
10-fold CV
Mean AUROC
0.542
0.531
0.550
Borzoi
CADD
CADD + Borzoi

Figure S4: **Related to Figure 4. (A)** Comparison of three different attribution methods (Input-gated gradient saliency, ISM, and ISM Shuffle) in a local sequence window centered on variant rs1905542 in the intron of the SHTN1 gene. Putative TF binding motifs are annotated at the bottom and aligned with the input sequence (HOCOMOCO v11 PWMs with lowest p-values as determined by Tomtom MEME). **(B)** Predicted RNA-seq coverage (GTEx tissue 'Whole Blood') for the WLS gene when inducing variant rs72670481. Measured coverage in 6 individuals with the reference allele and in hetero- or homozygous individuals for the alternative allele is displayed below the predictions, along with attribution scores computed in a local window centered on the variant. The attributions scores are calculated with respect to the log of sum of exon coverage for the WLS gene. **(C)** Predicted RNA-seq coverage (GTEx tissue 'Whole Blood') for variant rs3890144, along with measured coverage in 8 individuals with the reference allele and in 8 individuals who are either hetero- or homozygous for the alternative allele. Attribution scores in a local window centered on the variant, calculated with respect to EPHB4 exon coverage, are displayed at the bottom. **(D)** The top-1-accuracy (percentage of eGenes which are correctly predicted) by posterior probability (PP)-weighted Borzoi scores compared to PP-weighted nearest gene across 49 GTEx tissues. **(E)** Left: Precision vs recall when classifying singleton variants from common variation (AF $> 0.05$) from gnomAD. Right: Average AUROC (10-fold cross validation).

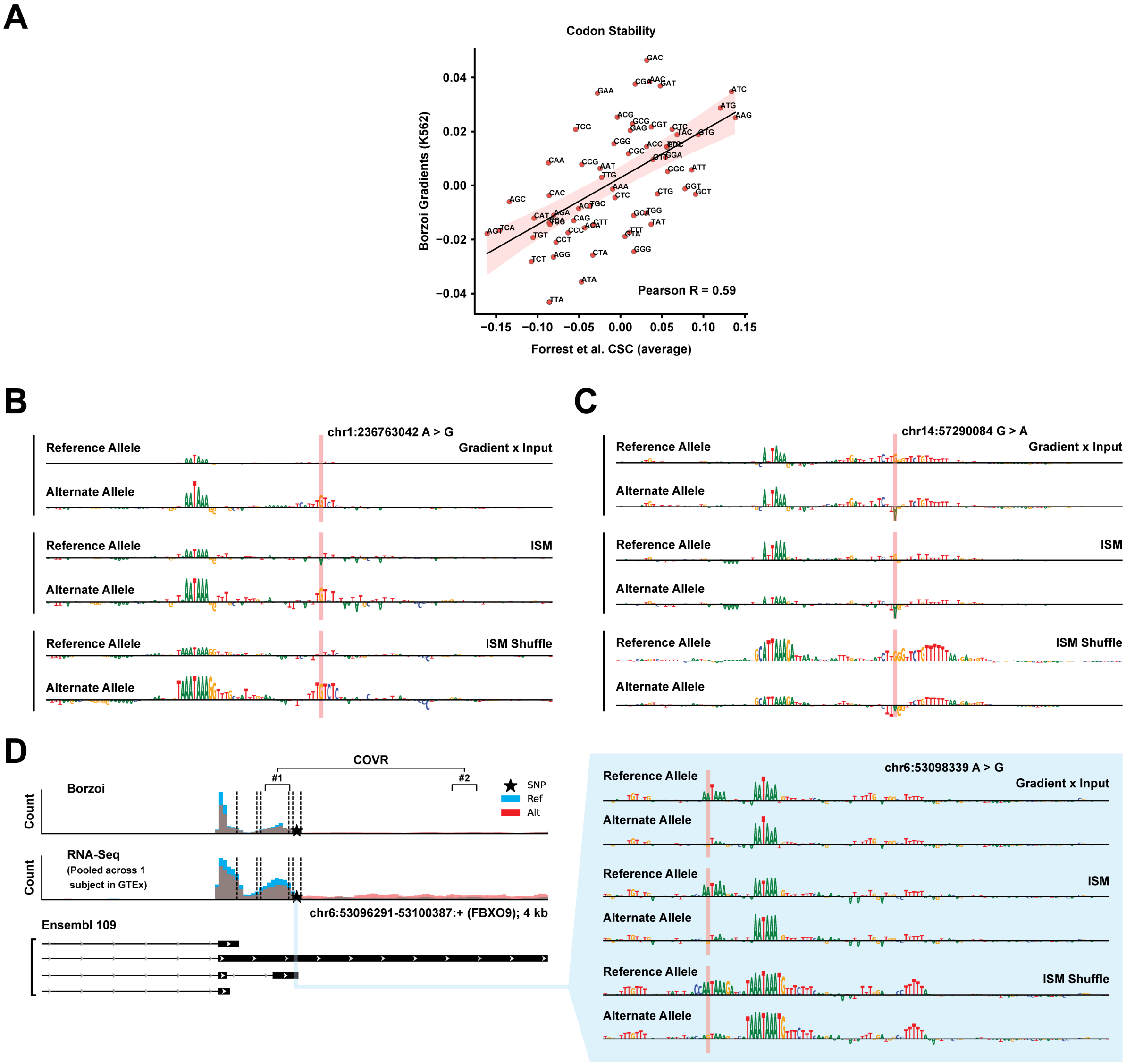


Figure S5: **Related to Figure 5.** **(A)** Comparison between the average gradient saliency of codons in genes from the Gasperini set and measured codon stability in data from Forrest et al. (2020). The gradient saliencies are calculated with respect to the predicted sum of exon coverage in K562 RNA-seq samples. **(B)** Attribution scores for a polyadenylation signal (PAS) in the ACTN2 gene when inducing variant rs114880747. The resulting scores of three separate attribution methods are displayed. The attribution scores are calculated with respect to the log ratio of coverage immediately upstream of the mutated PAS relative to coverage upstream of the distal-most PAS. (**C**) Attribution scores for a PAS in the AP5M1 gene when inducing variant rs80168986, using three attribution methods. **(D)** Predicted RNA-seq coverage (GTEx pooled) for variant rs74327114 (occurring in gene FBXO9), along with measured coverage in 1 individual with the reference allele and 1 heterozygous individual (two tissues each). Attribution scores, using three separate methods, of PAS coverage log ratio are displayed to the right and indicate loss of an extra hexamer motif, resulting in moderate reduction in polyadenylation efficiency.

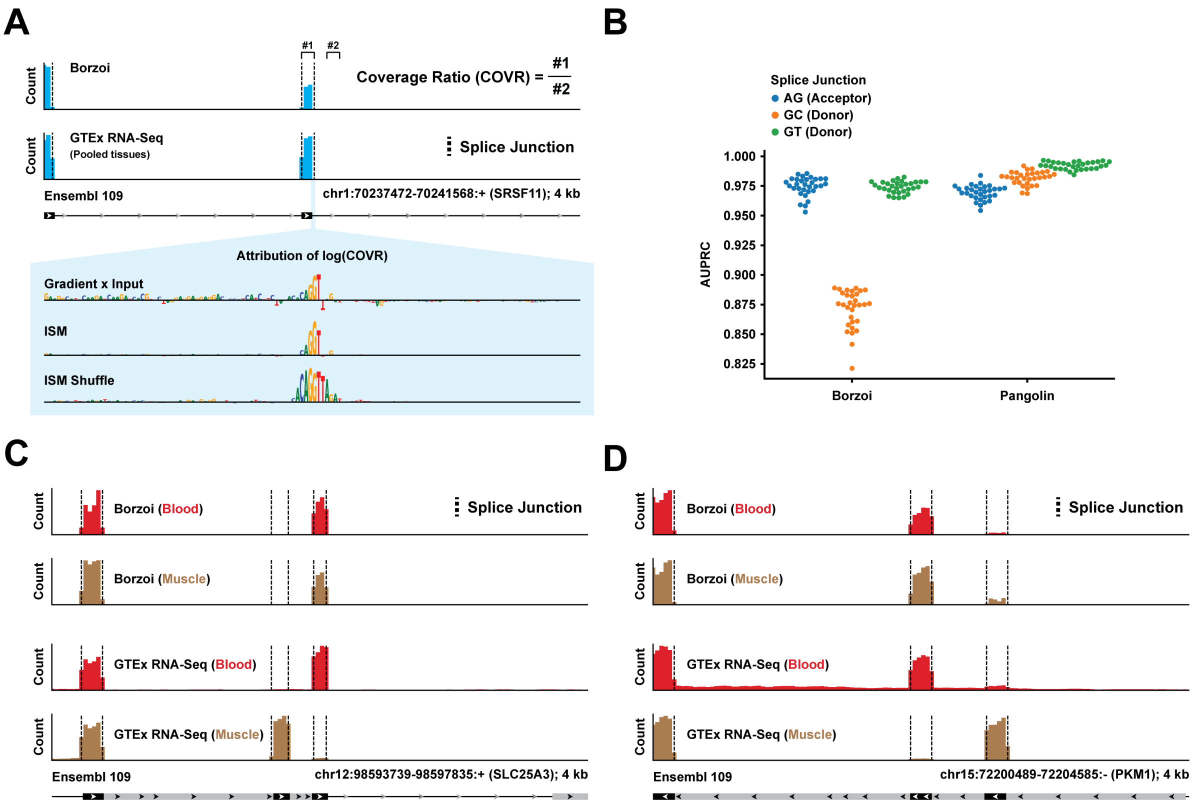


Figure S6: **Related to Figure 6.** **(A)** Predicted and measured RNA-seq coverage across an exon in the SRSF11 gene (GTEx pooled-tissue), centered on its splice donor. Attribution scores in a local window around the splice donor are displayed below the predicted tracks. The attribution scores are calculated with respect to the log ratio of exon-to-intron coverage. **(B)** Comparison between Borzoi's predicted exon-to-intron coverage ratio statistic and Pangolin's predicted splice usage when classifying annotated splice donors/acceptors from matched negatives in the reference genome. Average precision (AUPRC) is displayed separately for each type of splice junction (AG - Acceptor, GT/GC - Donor) and each dot corresponds to a GTEx tissue. **(C)** Predicted and measured RNA-seq coverage across an alternative splicing event in the SLC25A3 gene for GTEx tissues 'Whole Blood' and 'Muscle' (both predictions and measurements are pooled across 3 tissue-specific samples for blood and muscle). **(D)** Predicted and measured whole blood- and muscle RNA-seq coverage across an alternative splicing event in the PKM1 gene (Predictions and measurements are pooled across 3 samples per tissue).

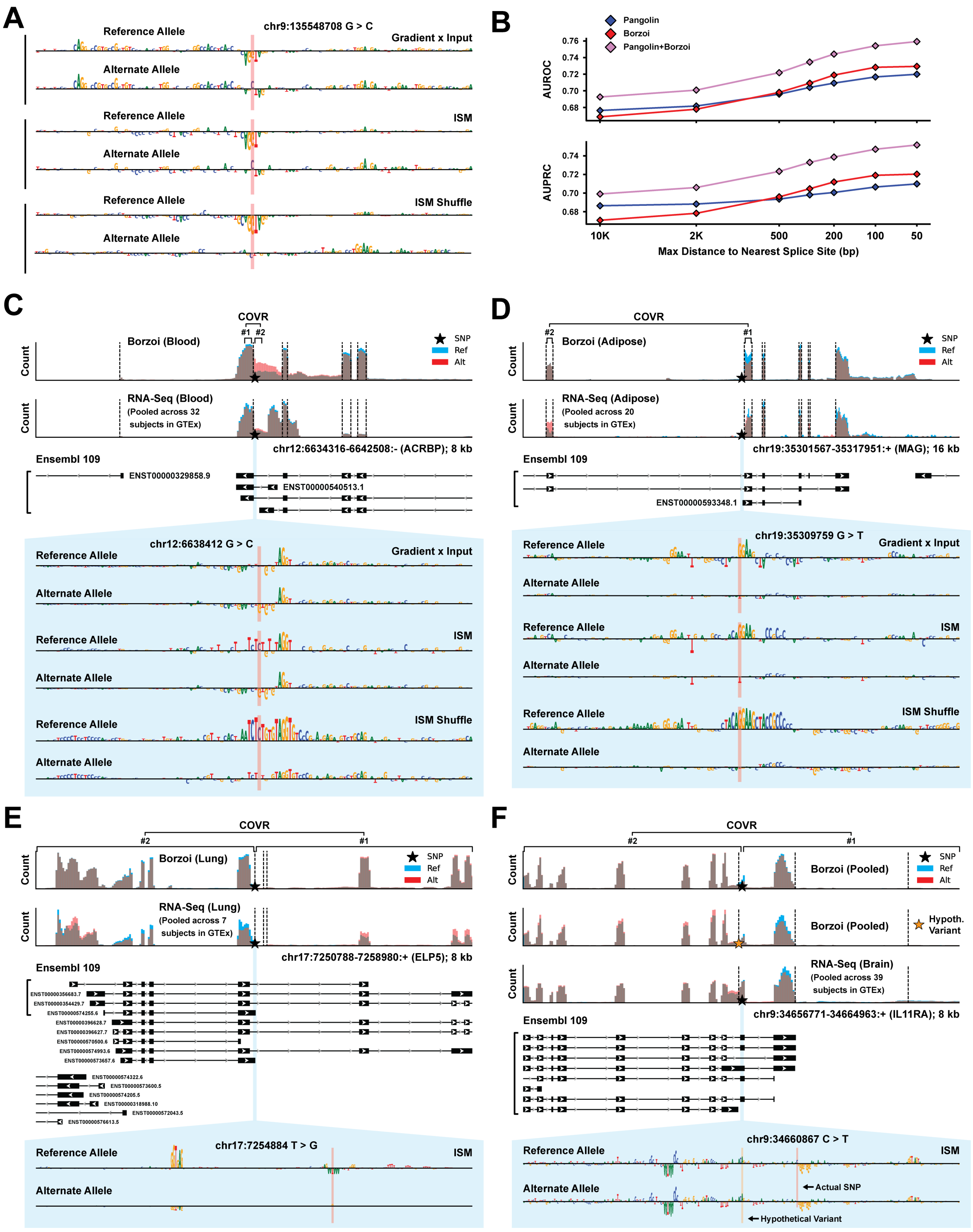
A
chr9:135548708 G > C
Reference Allele
Alternate Allele
Gradient x Input
ISM
ISM Shuffle
B
Pangolin
Borzoi
Pangolin+Borzoi
AUROC
AUPRC
10K
2K
500
200
100
50
Max Distance to Nearest Splice Site (bp)
C
COVR
#1
#2
Borzoi (Blood)
RNA-Seq (Blood)
(Pooled across 32 subjects in GTEx)
SNP
Ref
Alt
Count
chr12:6634316-6642508:- (ACRBP); 8 kb
Ensembl 109
ENST00000329858.9
ENST00000540513.1
chr12:6638412 G > C
D
Borzoi (Adipose)
RNA-Seq (Adipose)
(Pooled across 20 subjects in GTEx)
chr19:35301567-35317951:+ (MAG); 16 kb
ENST00000593348.1
chr19:35309759 G > T
E
Borzoi (Lung)
RNA-Seq (Lung)
(Pooled across 7 subjects in GTEx)
chr17:7250788-7258980:+ (ELP5); 8 kb
chr17:7254884 T > G
F
Borzoi (Pooled)
Hypoth. Variant
RNA-Seq (Brain)
(Pooled across 39 subjects in GTEx)
chr9:34656771-34664963:+ (IL11RA); 8 kb
chr9:34660867 C > T
Actual SNP
Hypothetical Variant

Figure S7: **Related to Figure 6.** **(A)** Attribution scores of exon-to-intron log coverage ratio in a local window centered on the variant rs55695858, comparing three separate methods. **(B)** Tissue-pooled variant effect prediction benchmark, comparing the performance of Pangolin, Borzoi and an ensemble of the two models (their average rank predictions) on the task of classifying between fine-mapped causal sQTLs (PP $> 0.9$) and expression- and distance-controlled negatives. A similar type of benchmark is presented in Figure 6D, but the results are separated by GTEx tissue in that analysis. **(C)** Predicted RNA-seq coverage (GTEx tissue 'Whole Blood') for variant rs1882553 using Borzoi, along with measured coverage in 32 individuals with the reference allele and 32 hetero- or homozygous individuals for the alternative allele (whole blood samples). Attribution scores (bottom) are computed with respect to the predicted log ratio of exon-to-intron coverage, comparing three methods (displayed in reverse-complemented form). **(D)** Predicted RNA-seq coverage (GTEx tissue 'Adipose') for variant rs10411704, along with measured coverage in 20 individuals with the reference allele and 20 hetero- or homozygous individuals for the alternative allele (adipose samples). Attribution scores of the predicted exon-to-exon log coverage ratio are displayed at the bottom, comparing three methods. **(E)** Predicted RNA-seq coverage (GTEx tissue 'Lung') and measured coverage in 7 individuals with the alternative (hetero- or homozygous) or reference allele of variant rs402514. Attribution scores (bottom) are calculated with respect to the log ratio of downstream-to-upstream coverage relative to the SNP. **(F)** Predicted RNA-seq coverage (GTEx pooled-tissue) and measured coverage in 39 individuals with the alternative (hetero- or homozygous) or reference allele of variant rs11575580. Attributions (bottom) are calculated with respect to the log ratio of downstream-to-upstream coverage.

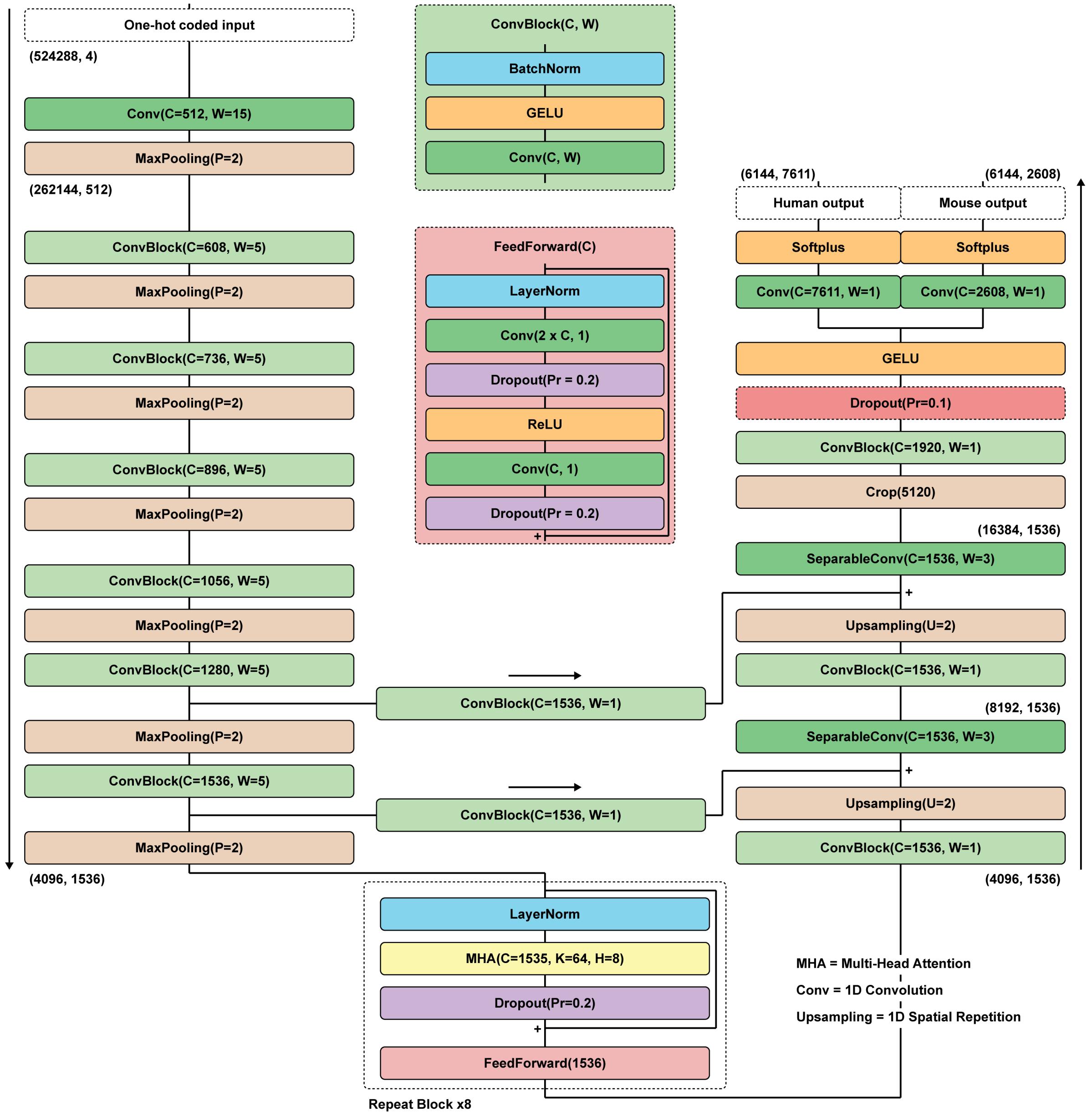


Figure S8: **Borzoi model architecture.** Neural network diagram. All convolutions are performed with a constant dilation rate of 1.
